# eIF4A2 targets developmental potency and histone H3.3 transcripts for translational control of stem cell pluripotency

**DOI:** 10.1101/2021.12.24.474136

**Authors:** Dan Li, Jihong Yang, Xin Huang, Hongwei Zhou, Jianlong Wang

**Author notes:** Corresponding Author: Jianlong Wang, PhD Professor, Department of Medicine, Columbia Center for Human Development (CCHD) Columbia Stem Cell Initiative (CSCI), Columbia University Irving Medical Center (CUIMC), William Black Building, 8^th^ Fl. Room 816, 650 W. 168^th^ St., New York, NY 10032, Office: BB8-801C.

## Abstract

Translational control has emerged as a fundamental regulatory layer of proteome complexity that governs cellular identity and functions. As initiation is the rate-limiting step of translation, we carried out an RNAi screen for key translation initiation factors required to maintain embryonic stem cell (ESC) identity. We identified eIF4A2 and defined its mechanistic action through Rps26-independent and -dependent ribosomes in translation initiation activation of mRNAs encoding pluripotency factors and the histone variant H3.3 with demonstrated roles in maintaining stem cell pluripotency. eIF4A2 also mediates translation initiation activation of Ddx6, which acts together with eIF4A2 to restrict the totipotent 2-cell transcription program in ESCs through Zscan4 mRNA degradation and translation repression. Accordingly, knockdown of eIF4A2 disrupts ESC proteome causing the loss of ESC identity. Collectively, we establish a translational paradigm of the protein synthesis of pluripotency transcription factors and epigenetic regulators imposed on their established roles in controlling pluripotency.

## Introduction

Cellular identity is driven by widespread gene expression control in multiple regulatory layers, with heterogeneity in the cellular epigenome, transcriptome, and proteome. Although initial work focused on dissecting the transcriptome and epigenome in safeguarding stem cell identity (*1*), RNA expression cannot directly determine protein abundance and cellular identity. Increasing studies have revealed the importance of post-transcriptional control in embryonic stem cells (ESCs) (*2*), and mRNA translation ranked first among all the enriched biological processes in analyzing the genes necessary for ESC maintenance (*3*). In ESCs, protein abundance and chromatin landscapes are susceptible to the alternations of translational control (*4*). However, mechanisms of mRNA translational control, particularly the rate-limiting translation initiation control in safeguarding ESC identity, remain poorly defined.

Mouse ESCs do not usually differentiate into extraembryonic trophoblast lineage, except for a minor population of bipotential 2C-like cells with both embryonic and extraembryonic differentiation propensities (*5*). While genetic manipulation of transcription programs and epigenetic machinery (*6-8*) can overcome this barrier, it is currently unknown whether a translational control mechanism exists to restrict the totipotent 2C program and extraembryonic lineage propensity in maintaining pluripotency.

Starting from an RNAi screen to identify key translation initiation factors (TIFs) that are required for maintaining ESC identity, we discovered in this study that eIF4A2 mediates a unique translation program by acting as both a translation activator and a repressor to control the expression of cellular potency regulators, including pluripotency factors and totipotency regulators, and epigenetic regulators, including a specific histone variant and a polycomb protein, which shapes the proteome of ESCs in safeguarding pluripotent stem cell identity.

## Results

### An RNAi screen identifies eIF4A2 as a critical TIF for ESC maintenance

TIFs include eukaryotic translation initiation factors (eIFs) and other factors involved in translation initiation (*9*) (Fig. 1A). While TIF RNA expression is dynamically regulated during mouse early embryogenesis, most TIF proteins are uniformly upregulated to the highest level at the blastocyst stage (*10*) (fig. S1A), from which ESCs can be derived, suggesting a potentially profound translation initiation control in ESCs.

**Fig. 1:**
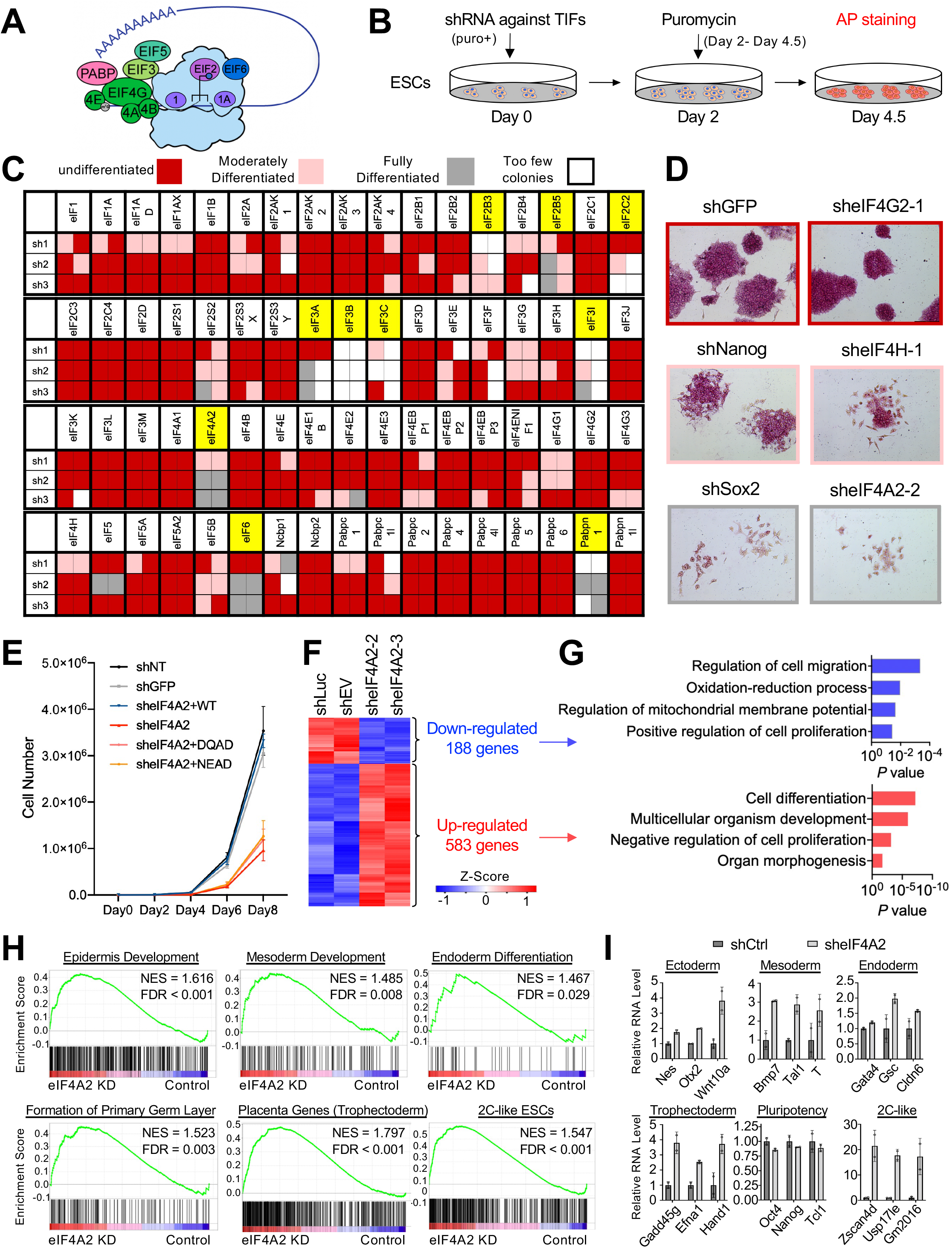
An RNAi screen identifies the requirement of eIF4A2 for maintaining the ESC identity. (**A**) Schematic of the eukaryotic cap-dependent translation initiation process. (**B**) Schematic of the RNAi screen to identify translation initiation factor dependency in ESCs. Puro+: The shRNA plasmid contains a puromycin-resistant gene. AP is alkaline phosphatase. (**C**) The RNAi screen results. The box colors denote ESC states as indicated. The selected candidates are highlighted in yellow. sh1-3 are three short hairpins for each gene, and screening was performed in biological replicates. (**D**) Representative examples of the AP-stained colony results from the RNAi screen, including the positive controls (shNanog, shSox2) and the negative control (shGFP). The border colors match the box colors used in (**C**). (**E**) Proliferation curves for ESCs with control KD [shNT (shNon-Targeting), shGFP], eIF4A2 KD (sheIF4A2), eIF4A2 KD rescued with WT eIF4A2 (sheIF4A2+WT) or helicase-activity mutants of eIF4A2 (sheIF4A2+DQAD/NEAD). (**F** and **G**) Heatmap (**F**) and Gene Ontology (GO) analysis (**G**) of the upregulated and downregulated transcripts upon eIF4A2 KD from RNA-seq data. shLuc: shLuciferase. EV: empty vector control. (**H**) GSEA results of primary germ layer gene sets (epidermis development, mesoderm development, endoderm differentiation, and formation of primary germ layer), placenta (trophectoderm) gene set, and 2C-like ESCs gene set (Z4 event: Zscan4 expression) by comparing eIF4A2 KD with control KD cells from RNA-seq data. NES, normalized enrichment score. (I) Examples of the RNA-seq result in multiple groups. shCtrl (control shRNA) includes shLuc and shEV control KD experiments.

To explore the functional significance of TIFs in regulating ESC identity, we performed an RNAi screen with three independent and constitutive short-hairpin RNAs (shRNAs; sh1-3 in Fig. 1C) targeting all 68 TIFs for ESC maintenance (Fig. 1, B and C). The screen was repeated once with high reproducibility (the two columns in Fig. 1C) and resulted in various ESC statuses ranging from undifferentiated (red; shGFP/sheIF4G2 as control/hit example) to moderately differentiated (pink; shNanog/sheIF4H as control/hit example) to fully differentiated (grey; shSox2/sheIF4A2 as control/hit example) cell states (Fig. 1, C and D). Candidate hits that resulted in cell death/loss were scored as “too few colonies” (white) (Fig. 1C). We found that knockdown (KD) of the TIFs previously reported for ESC maintenance (*3, 11*), such as eIF2B3 and eIF2S2, also showed moderate differentiation (Fig. 1C), and that depletion of eIF4G2 (aka Nat1) maintains, or even slightly enhances, typical dome-shaped and alkaline phosphatase (AP)-positive ESC morphology as reported (*12*) (Fig. 1, C and D), supporting the validity of our screen. Overall, our screen of 68 TIFs identified 10 TIFs (Fig. 1C, highlighted in yellow) whose depletion induced severe differentiation and/or death in ESCs. To identify TIFs specifically required for ESC pluripotency instead of cellular viability, we removed eight fitness genes essential to many immortalized and cancer cell types (*13*) (fig. S1B) and one gene likely required for ESC viability (eIF2C2, Fig. 1C). eIF4A2 is the only and most consistent TIF whose depletion causes moderate to severe ESC differentiation instead of cell death (Fig. 1, C and D, and fig. S1C), which is consistent with its specific expression enrichment in preimplantation inner cell mass (ICM) of the blastocyst in vivo (fig. S1D) and its downregulated expression during RA (retinoic acid) or FA (Fgf2/Activin A) differentiation in vitro (fig. S1, E and F).

Next, we validated the KD efficiency (fig. S1, G, H and K) and the differentiation phenotype in an alternative ESC line (fig. S1I) and under an alternative pluripotent state of 2i/LIF (*14*) cultured naïve ESCs (fig. S1, J and K). eIF4A2 KD had a minimal effect on cell death/apoptosis of non-pluripotent NIH/3T3 cells and mouse embryonic fibroblasts (MEFs) (fig. S1, L-N), as well as pluripotent cells (fig. S1O). While eIF4A2 depletion didn’t affect proliferation of non-pluripotent cells, eIF4A2 KD decreased proliferation of pluripotent ESCs (Fig. 1E). And only WT, but not helicase activity dead mutants (DQAD and NEAD) of eIF4A2 (*15*), can rescue the eIF4A2 KD effects on ESC proliferation, morphology, and the expression of differentiation and 2C transcripts (Fig. 1E, fig. S1, P and Q). These results established the essential role of eIF4A2 in maintaining the pluripotency. However, overexpression of eIF4A2 in ESCs (fig. S2A) had minimal effects on the ESC morphology, proliferation, or expression levels of pluripotency factors (fig. S2, A-D), suggesting eIF4A2 is not limiting in ESC maintenance. To explore its potential roles in establishing the pluripotency, we study the loss/gain of eIF4A2 in reprogramming of somatic cells to induced pluripotent stem cells (iPSCs). Despite its dispensability in MEFs (fig. S1, M and N), eIF4A2 KD dramatically decreased MEF reprogramming efficiency by Yamanaka factors (*16*) (fig. S2, E to G) and also decreased reprogramming efficiency of pre-iPSCs to iPSCs (fig. S2, H to J). However, eIF4A2 overexpression had minimal effects on MEF or pre-iPSC reprogramming efficiencies (fig. S2, E to J). These results together establish the critical roles of eIF4A2 in the maintenance and establishment of stem cell pluripotency.

To characterize the molecular features of eIF4A2-depleted ESCs, we performed RNA-seq of ESCs treated with control shRNAs (shCtrl) against empty vector (shEV) or luciferase (shLuc) and shRNAs against eIF4A2 (sheIF4A2) (using the same KD and drug selection time points as those for the TIF screen in Fig. 1B) (table S1). Gene ontology (GO) analysis of RNA-seq data of 188 downregulated genes and 583 upregulated genes (Fig. 1F) revealed the enrichment of GO terms “positive regulation of cell proliferation” and “negative regulation of cell proliferation”, respectively (Fig. 1G), consistent with the compromised growth of eIF4A2 KD ESCs (Fig. 1E). Of note, this proliferation defect of KD ESCs is also associated with increased “cell differentiation”, a GO term in the upregulated gene list (Fig. 1, F and G) without changes in cell death or apoptosis (fig. S1O), suggesting that differentiation is likely the cause for reduced growth of eIF4A2 KD ESCs. Indeed, gene set enrichment analysis (GSEA) revealed the enrichment of gene sets representing all primary germ layers, and surprisingly, placenta (trophectoderm) transcripts as well as Zscan4 event-associated 2C-like ESC transcripts (*17*) in eIF4A2 KD ESCs (Fig. 1H). Despite the upregulation of the RNA levels of three primary germ layers, trophectoderm, and 2C genes, eIF4A2 KD didn’t change the RNA levels of master pluripotency genes (Fig. 1I). These results highlight the profound role of eIF4A2 in restricting early developmental potency and embryonic/extraembryonic differentiation potential for ESC maintenance.

### eIF4A2 is responsible for translation activation and repression of mRNAs encoding cellular potency factors

eIF4A2 is a translation initiation factor, but its depletion did not alter global translation rates in ESCs (Fig. 2A). The lack of a global effect on translation upon eIF4A2 loss was also reported in non-pluripotent NIH/3T3, HeLa, and HEK293 cell lines (*18, 19*). These results suggest that eIF4A2 likely acts on a specific set of mRNAs for the translation initiation control of stem cell pluripotency.

**Fig. 2:**
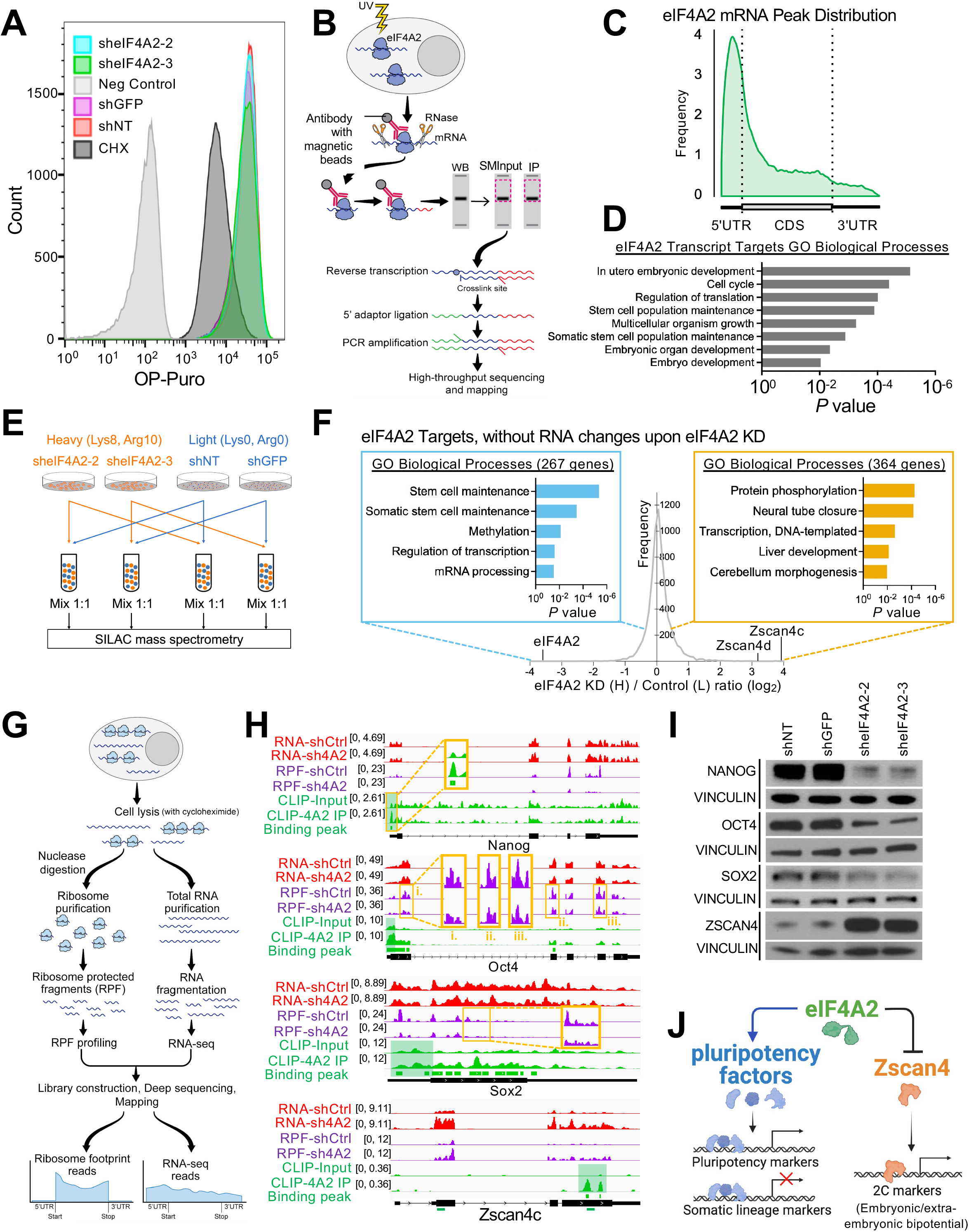
eIF4A2 acts as both a translation activator and repressor of specific mRNAs encoding cellular potency factors. (**A**) Flow cytometry for OP-puro (O-Propargyl-puromycin) incorporation in ESCs with control KD and eIF4A2 KD. OP-puro is used to label nascent peptides, indicating a global translation level. CHX, cycloheximide, a protein synthesis inhibitor. (**B**) Schematic diagram of the eCLIP-seq protocol. (**C**) Peak distribution around mRNAs of eIF4A2 targets, identified by eCLIP-seq. (**D**) GO analysis of eIF4A2 targets identified by eCLIP-seq. (**E**) Schematic diagram of SILAC-MS experiment protocol (details in Methods). (**F**) Graph of the frequency distribution of heavy/light (H/L, eIF4A2 KD/control) ratios of all proteins identified by SILAC-MS, with GO analysis of the genes which are targeted by eIF4A2 with decreased (left side, blue box) or increased (right side, orange box) protein levels while maintained mRNA levels upon eIF4A2 KD. (**G**) Schematic diagram of the ribosome profiling protocol. (**H**) IGV (Integrative Genomics Viewer) snapshots on Nanog, Oct4, Sox2, and Zscan4 showing: top (red) as RNA-seq data between control KD and eIF4A2 KD; middle (purple) as RPF profiling data between control KD and eIF4A2 KD; bottom (green) as eIF4A2 binding (eCLIP-seq) profiling data, input as control, binding of eIF4A2 at TIRs (for Nanog, Oct4, and Sox2) and the CDS of Zscan4c near 3’ UTR is shaded in green. shCtrl: shRNA control; sh4A2: sheIF4A2. The green marks below the Zscan4c gene architecture diagram are the CLIP-qPCR amplicon positions in Fig. 6F. (**I**) Western blots of NANOG, OCT4, SOX2, and ZSCAN4 in ESCs with control KD and eIF4A2 KD. VINCULIN serves as the loading control. (**J**) A model depicting eIF4A2-mediated translation activation of pluripotency transcripts and repression of Zscan4 in ESCs.

To identify specific mRNAs whose translation initiation is regulated by eIF4A2 in ESCs, we first identified direct RNA binding targets of eIF4A2 by performing enhanced UV crosslinking immunoprecipitation coupled with high-throughput sequencing (eCLIP-seq) (*20*) (Fig. 2B) with biological replicates, which yielded reproducible results (fig. S3A). The binding peaks of eIF4A2 are enriched in mRNAs, particularly at 5’ UTR and the immediate neighboring region of CDS, an mRNA translation initiation region (TIR) that is crucial for the regulation of translation initiation (*21*) (Fig. 2C, and fig. S3, B and C, and table S2). The GO analysis of the enriched binding sites revealed that its targets are important for embryonic development, stem cell maintenance, and regulation of translation (Fig. 2D).

Second, we explored the proteome changes upon eIF4A2 KD using SILAC (stable isotope labeling by amino acids in cell culture)-based quantitative MS (Fig. 2E). Upon eIF4A2 KD, the downregulated proteins were enriched for amino acid biosynthetic process and stem cell maintenance, whereas the upregulated proteins were significantly clustered into protein phosphorylation and development-related processes (fig. S3D and table S2). We filtered these proteins with maintained mRNA abundance upon eIF4A2 depletion and only chose the eIF4A2-binding targets, obtaining a list of 267 and 364 proteins that are decreased and increased, respectively, in eIF4A2 KD relative to control KD ESCs (Fig. 2F, and fig. S3E, and table S2). These targets were considered as directly subject to the translation initiation control by eIF4A2. Notably, the GO analysis of the downregulated proteins showed the enrichment of stem cell maintenance as the top term (Fig. 2F), among which are 16 pluripotency-associated factors, including the core transcription factors Nanog, Oct4, and Sox2, and all of them were targeted by eIF4A2 at their TIRs (fig. S3E and table S2). In contrast, the top upregulated proteins upon eIF4A2 KD were Zscan4c/d (Fig. 2F). This well-known 2C factor can activate its own transcription and the associated 2C molecular program (*17, 22*), consistent with our GSEA result (Fig. 1H).

Third, we performed ribosome profiling (Fig. 2G) to validate the translational outcome of the above-identified eIF4A2 targets. The ribosome profiling results showed that eIF4A2 KD didn’t change the ribosome bindings on the housekeeping gene Vcl mRNA (encoding VINCULIN) or the key global translation control gene Mtor mRNA (fig. S3F), confirming that eIF4A2 KD didn’t change the global translation (Fig. 2A). We then confirmed that a number of pluripotency regulators, including the core factors Nanog, Oct4, and Sox2, are translationally activated by eIF4A2 with maintained mRNA levels but decreased binding of translating ribosomes (RPFs, ribosome protected mRNA fragments) upon eIF4A2 KD (Fig. 2H and fig. S3G). The binding of eIF4A2 at TIRs (shaded in green in Fig. 2H and fig. S3G) and decreased protein levels of these pluripotency factors after eIF4A2 KD (Fig. 2I) were further confirmed. We also confirmed that eIF4A2 KD increased the RNA and RPF levels of Zscan4c/d, the ZSCAN4 protein levels, and ZSCAN4-positive cells in ESC culture (Fig. 2, H and I, and fig. S3, G and H). Of note, eIF4A2 binds to Zscan4c/d mRNAs at the CDS (coding sequence) near 3’ UTRs, instead of TIRs (Fig. 2H and fig. S3G).

These results demonstrate that eIF4A2 activates translation initiation of pluripotency transcripts and represses the expression of the totipotency 2C marker Zscan4, which in turn restricts embryonic and extraembryonic lineage propensities in ESCs (*23*) (Fig. 1H and 2J).

### eIF4A2 depletion impacts ribosome binding at translation initiation regions

To explore the mechanism underlying eIF4A2-mediated translation control, we compared genome-wide transcriptional and translational differences upon eIF4A2 depletion. Consistent with observations before, the homo-directional upregulated genes are enriched with differentiation genes (including trophectoderm markers Gcm1 and Fgfr2) as well as 2C markers (e.g., Zscan4c/d), and the homo-directional downregulated genes are enriched for genes involved in negative regulation of mitochondrial fusion, which is also associated with cell differentiation (*24*) (Fig. 3, A top and B, and fig. S4, A and B, and table S3). Notably, upon eIF4A2 depletion, the distribution of ribosome binding is shifted, and the enrichment at TIR observed in the control is lost (Fig. 3C), indicating that eIF4A2 KD compromised the ribosome binding at TIR with a greater extent than the ribosome binding on the other regions of mRNAs. Indeed, the comparison of transcriptome and translation initiation “regulome” showed that more expression changes upon eIF4A2 KD occurred on the ribosome binding density at TIR (TIR_RPF, Y-axis), instead of RNA levels (X-axis) (Fig. 3A middle and table S3), and there are more changes happening on TIR_RPF (Fig. 3A middle) than RPF (Fig. 3A top).

**Fig. 3:**
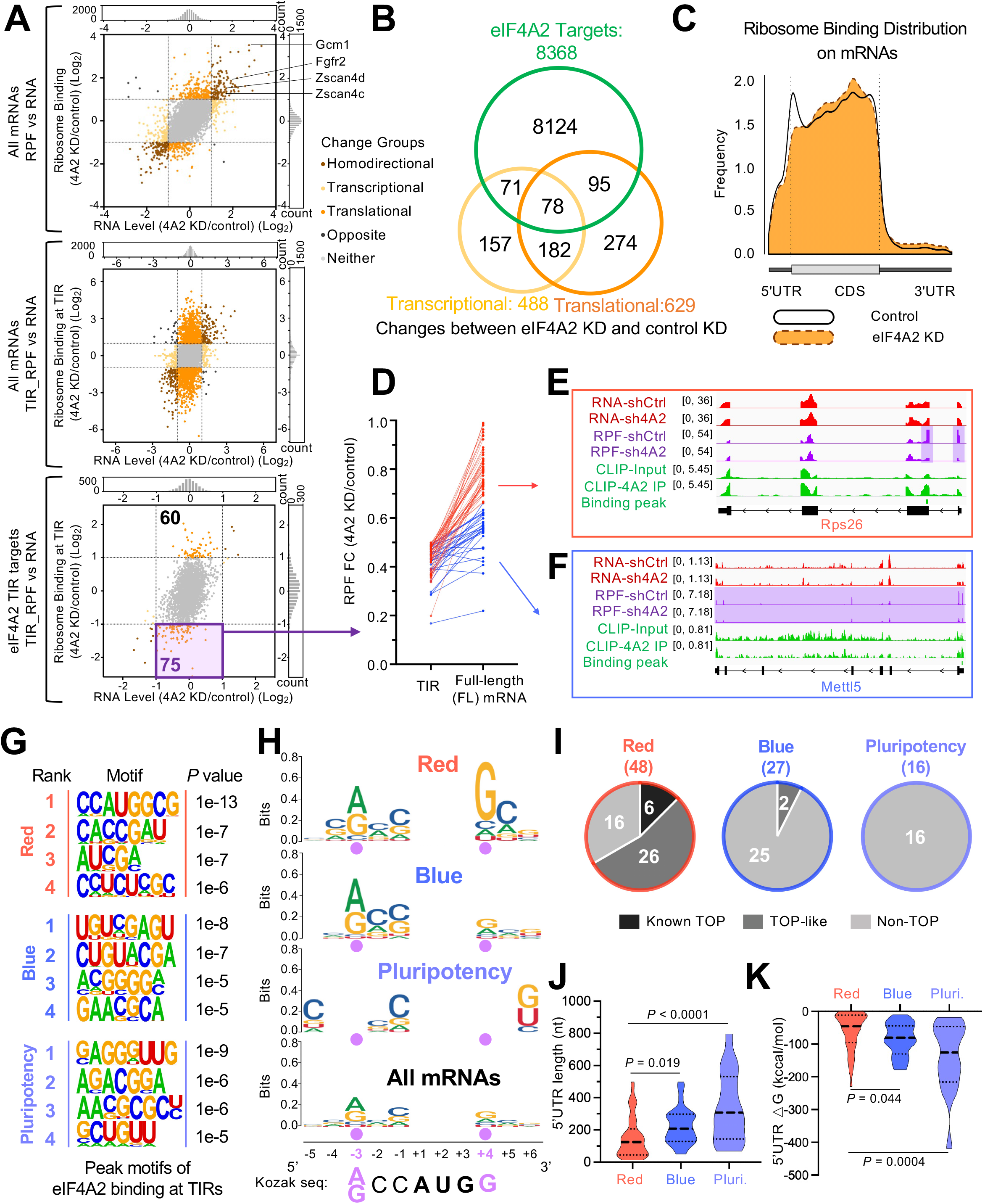
eIF4A2 activates target mRNAs in two modes of action depending on their 5’ UTR complexity and the Kozak consensus at TIRs. (**A**) Comparing eIF4A2 KD with control KD ESCs, top: transcriptional and translational changes of all mRNAs; middle: transcriptional and translation initiation changes of all mRNAs; bottom: transcriptional and translation initiation changes of eIF4A2 TIR targets. Targets (75) with ribosome binding at TIR decreased upon eIF4A2 KD are indicated. (**B**) Venn diagram showing mRNAs with transcriptional changes, translational changes, and with eIF4A2 binding targets. (**C**) Ribosome binding distribution around mRNAs, comparing eIF4A2 KD and control, identified by ribosome profiling. (**D**) eIF4A2 activates the translation initiation of two distinct groups. Each line represents one target from the purple box in (A). The left and right ends of the line represent RPF FC between eIF4A2 KD and control KD at TIR and full-length (FL) mRNA body, respectively. (**E** and **F**) IGV snapshots of representative red target Rps26 (**E**) and representative blue target Mettl5 (**F**) with the same datasets as shown in Fig. 2H. (**G**) Enriched motifs in eIF4A2 bound peaks at TIRs of the red (top), blue (middle), and pluripotency (bottom) targets. (**H**) Sequence conservation analysis of red, blue, and pluripotency targets around the start codons relative to the Kozak sequence. The two critical consensus nucleotides (−3 and +4) within the Kozak sequence are highlighted with purple dots. (**I**) Pie charts showing the red, blue, and pluripotency targets with known 5’ TOP (5’ terminal oligopyrimidine tract), TOP-like, or non-TOP mRNAs. (**J** and **K**) Graphs showing 5’ UTR length (**J**) and free unfolding energy (kcal/mol) (**K**) of the red, blue, and pluripotency (pluri.) targets.

To identify the candidates through which eIF4A2 exerts the translation initiation control, we applied the stringent criteria to filter the eIF4A2 targets (see details in Methods). To focus on the molecular mechanisms underlying the ribosome recruitment mediated by eIF4A2, we only focus on the top candidates with the most significant ribosome binding changes at TIRs upon eIF4A2 depletion. These consist of 60 and 75 genes with translation initiation increase and decrease, respectively, upon eIF4A2 KD (Fig. 3A bottom and table S4). Among the 60 upregulated genes, many genes can induce ESC differentiation or are important for cellular development, such as Hexim1 (*25*), Tcf12 (*26*), Tcf3 (*27*), and Tet1 (*28*). Thus, in eIF4A2-depleted ESCs, the overexpression of these genes (at protein level) may contribute to the loss of pluripotency and cellular differentiation. Conversely, among the 75 downregulated genes, eIF4A2 KD eliminated the ribosome binding at TIRs, observed in the control KD, of many pluripotency regulators (Fig. 2H and fig. S3G). The pluripotency program was thus directly downregulated through the loss of the translation initiation activation upon eIF4A2 depletion. These results indicate the importance of the translation initiation activation exerted by eIF4A2 in maintaining ESC identity. We, therefore, focused on characterizing the 75 genes whose translation initiation was activated by eIF4A2 to explore how eIF4A2 exerts translation initiation activation control.

### eIF4A2 activates target mRNAs via Kozak context-dependent and independent translation initiation

We first examined the ribosome density changes at both TIR and full-length (FL) mRNA bodies of those 75 genes, revealing two distinct patterns of RPF’s decrease upon eIF4A2 KD (Fig. 3, D to F): 1) the RPF reduction mainly at TIRs without alteration on the rest of mRNA bodies (temporarily defined as “red targets”, red lines in Fig. 3D; Rps26 as one example in Fig. 3E); 2) the RPF reduction along FL mRNA bodies encompassing both TIRs and the rest of mRNA bodies (temporarily defined as “blue targets”, blue lines in Fig. 3D; Mettl5 as one example in Fig. 3F).

Translation regulation can be driven by functional RNA regulons within TIRs, such as RNA elements and structures (*21*). To understand how eIF4A2 may distinguish the red from blue targets in translation initiation activation, we scanned eIF4A2 binding motifs at TIRs of these targets (Fig. 3G). We found that the Kozak sequence, which functions as the translation initiation site mediating ribosome assembly (*21*), was enriched only in the red group (rank 1 in red; Fig. 3G). Analysis of the sequence enrichment around the start codons revealed that the Kozak consensus is more robust in the red group than in the blue group, considering the consensus with the most critical two nucleotide positions (G at +4, A/G at -3 relative to +1 as the beginning of the start codon) for the Kozak sequence (Fig. 3H). We also noticed a motif with a pyrimidine (C/U) stretch in the eIF4A2 binding motifs at red TIRs (rank 4 in the red; Fig. 3G), a prominent feature of the 5’ *t*erminal *<u>o</u>*ligo*<u>p</u>*yrimidine motif (5’ TOP) (*29*). 5’ TOP mRNAs encode various components of mRNA translation machinery, including ribosomal proteins, translation initiation/elongation factors, and some other proteins (*29*). We found six known 5’ TOP mRNAs (eIF3K, Hint1, Mrps21, Plp2, Rps26, and Rpl15) in the red group and none in the blue group (Fig. 3I). While examining the transcriptional start sites (TSSs) of the rest of mRNAs by utilizing the database of TSS (dbTSS) (*30*) and the Refseq, UCSC resources (fig. S4C), we found 26 red but only two blue mRNAs having TOP-like motifs (*31*) (Fig. 3I). Furthermore, compared with the red targets, the blue targets have longer 5’ UTR and more complicated RNA secondary structures, resulting in greater minimum free energy estimates (*21*) (Fig. 3, J and K, and fig. S4, D to F).

Of note, pluripotency-associated mRNA targets discussed before are considered as blue targets based on their RPF change patterns (Fig. 2H and fig. S3G) and the following 5’ UTR characteristics: like blue targets, pluripotency mRNAs bear weak Kozak consensus sequence (Fig. 3H), no known 5’ TOP mRNAs or TOP-like motifs (Fig. 3I), longer 5’ UTR (Fig. 3J), greater free energy estimates of 5’ UTR (Fig. 3K), and more complicated 5’ UTR structures (fig. S4, D to F).

Together, these results indicate that eIF4A2 activates translation initiation of two distinct groups of target mRNAs (red versus blue) depending on their 5’ UTR length/sequences/structures and the Kozak consensus at TIRs. RNA regulons can confer ribosome specificity to gene regulation (*21, 32*). The mRNAs with stringent Kozak sequences and short 5’ UTR are known to be encoded by RPS26-containing ribosomes, whereas RPS26-deficient ribosomes enhance translation of mRNAs with weak Kozak sequences and long 5’ UTR, clustering in specific regulatory pathways (*33*). We thus hypothesized that RPS26’s presence and absence in ribosomes might be responsible for differential translation initiation of eIF4A2-regulated red and blue targets, respectively.

### Rps26-dependent and -independent translation control can distinguish two modes of translation activation exerted by eIF4A2

To validate the identified features (stringent Kozak sequence and 5’ TOP motif) in conferring Rps26-dependent and -independent eIF4A2 functions, we first tested our hypothesis that the pluripotency target mRNAs are preferred by Rps26-deficient ribosomes. Supporting this, ESCs with Rps26 depletion through shRNA-mediated KD maintained both the typical dome-shaped ESC morphology (Fig. 4, A and B) and protein expression levels of the pluripotency targets, such as KLF5, OCT4, and SOX2, and even a slight increase of NANOG protein level (Fig. 4C). In contrast, Rps26 depletion caused the reduced protein abundances (Fig. 4C), but not RNA levels (Fig. 4D), of red targets H3.3 and Suz12, confirming the Rps26-dependent translation of red targets.

**Fig. 4:**
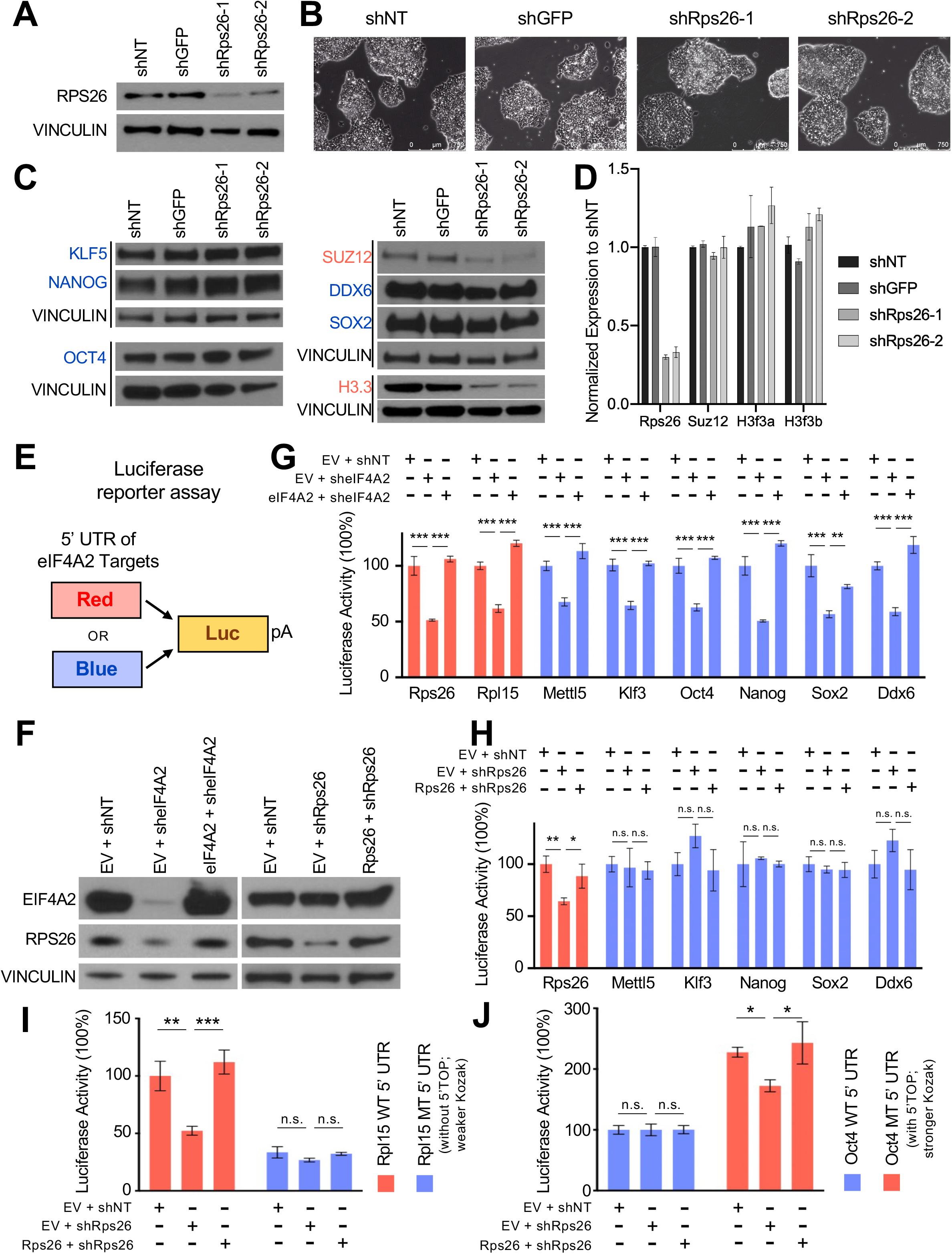
The two modes of translation activation exerted by eIF4A2 are in Rps26-dependent and -independent control. (**A** and **C**) Western blots of the indicated proteins in ESCs with control KD and Rps26 KD. (**B**) Cell morphology of ESCs with control KD and Rps26 KD. (**D**) the qRT-PCR result of Rps26, Suz12, H3f3a, and H3f3b in ESCs with control KD and Rps26 KD, relative to the expression levels in shNT and normalized to β-actin expression. (**E**) Schematic of the luciferase reporter assay. (**F**) Western blots of the indicated proteins in ESCs expressing the indicated shRNA and plasmid. (**G** and **H**) Luciferase activity in cells transfected with mRNAs containing the 5’ UTR of the red or blue target and expressed with the indicated shRNA and plasmid. (**I** and **J**) Luciferase activity in cells transfected with mRNAs containing the WT or mutated 5’ UTR of the Rpl15 (I) or Oct4 (J) and expressed with the indicated shRNA and plasmid.

We then employed the luciferase reporter assays to confirm that 5’ UTRs of red and blue targets are functionally effective in mediating translation initiation activation of corresponding luciferase reporter activities, demonstrated by downregulated luciferase activities upon eIF4A2 depletion, which can be rescued by sheIF4A2-immune eIF4A2 cDNA (Fig. 4, E to G). We further confirmed Rps26-dependent and -independent functions of eIF4A2 in differential translation control of red and blue targets by showing that the 5’ UTR of the red targets, but not the blue targets, downregulated luciferase activities upon Rps26 depletion, which can be rescued by shRps26-immune Rps26 cDNA (Fig. 4, E, F, and H). Of note, the shRps26 targets the CDS of Rps26 mRNA instead of 5’ UTR, so the shRps26 doesn’t target the Rps26 5’ UTR-luciferase reporter.

Finally, we asked whether a minimal alteration (swap) of 5’ UTR sequences between red and blue targets would correspondingly alter their translational response to the presence/absence of Rps26. By mutating the red target Rpl15 5’ UTR with the deletion of its 5’ TOP motif and disruption of the Kozak (fig. S5A), we found that the initially responsive and translationally rescuable Rpl15 5’ UTR became non-responsive to Rps26 KD or the transgenic rescue with shRps26-immune Rps26 cDNA (Fig. 4I). Conversely, by mutating the blue target Oct4 5’ UTR with the adoption of 5’ TOP from the red target Rpl15 5’ UTR and a single nucleotide change that created an optimized Kozak motif (fig. S5B), we found that the initially non-responsive Oct4 5’ UTR became responsive, i.e., downregulated, to Rps26 KD, which can be translationally rescued by shRps26-immune Rps26 cDNA (Fig. 4J). We also confirmed that these mutations didn’t affect the RNA stability (fig. S5, C and D).

Together, these results showed that eIF4A2 activates translation initiation of its targets in two modes of action, through Rps26-dependent and -independent ribosomes, by recognizing the characteristic features of 5’ UTRs, including Kozak sequence and 5’ TOP motif. Hereafter, we will mention these red and blue targets as Rps26-dependent and Rps26-independent targets, respectively.

### eIF4A2 activates the translation of H3.3 to repress trophectoderm development

We next examined the functional relevance of eIF4A2-mediated translation initiation of the Rps26-dependent targets in maintaining the ESC identity. GO analysis of these targets (table S4) revealed the predominant presence of H3.3-coding H3f3a and H3f3b genes in 8 out of 10 top terms (fig. S6A). Furthermore, supporting Rps26-dependent translation initiation activation of H3.3 by eIF4A2, depletion of Rps26 (Fig. 4, C and D) and eIF4A2 (Fig. 5, A and B, and fig. S6B) led to downregulation of H3.3 TIR_RPF/protein but not mRNA levels, and the overexpression of Rps26 in eIF4A2 KD cells could rescue the H3.3 protein level partially (Fig. 5, C and D). Noteworthily, such eIF4A2-mediated translation initiation activation is highly specific to H3.3, but not H3.1/H3.2, as eIF4A2 KD affects neither ribosome binding on H3.1/H3.2 mRNAs nor H3.1/H3.2 protein synthesis, even though eIF4A2 binds to H3.1/H3.2 mRNAs (Fig. 5B and fig. S6, C and D).

**Fig. 5:**
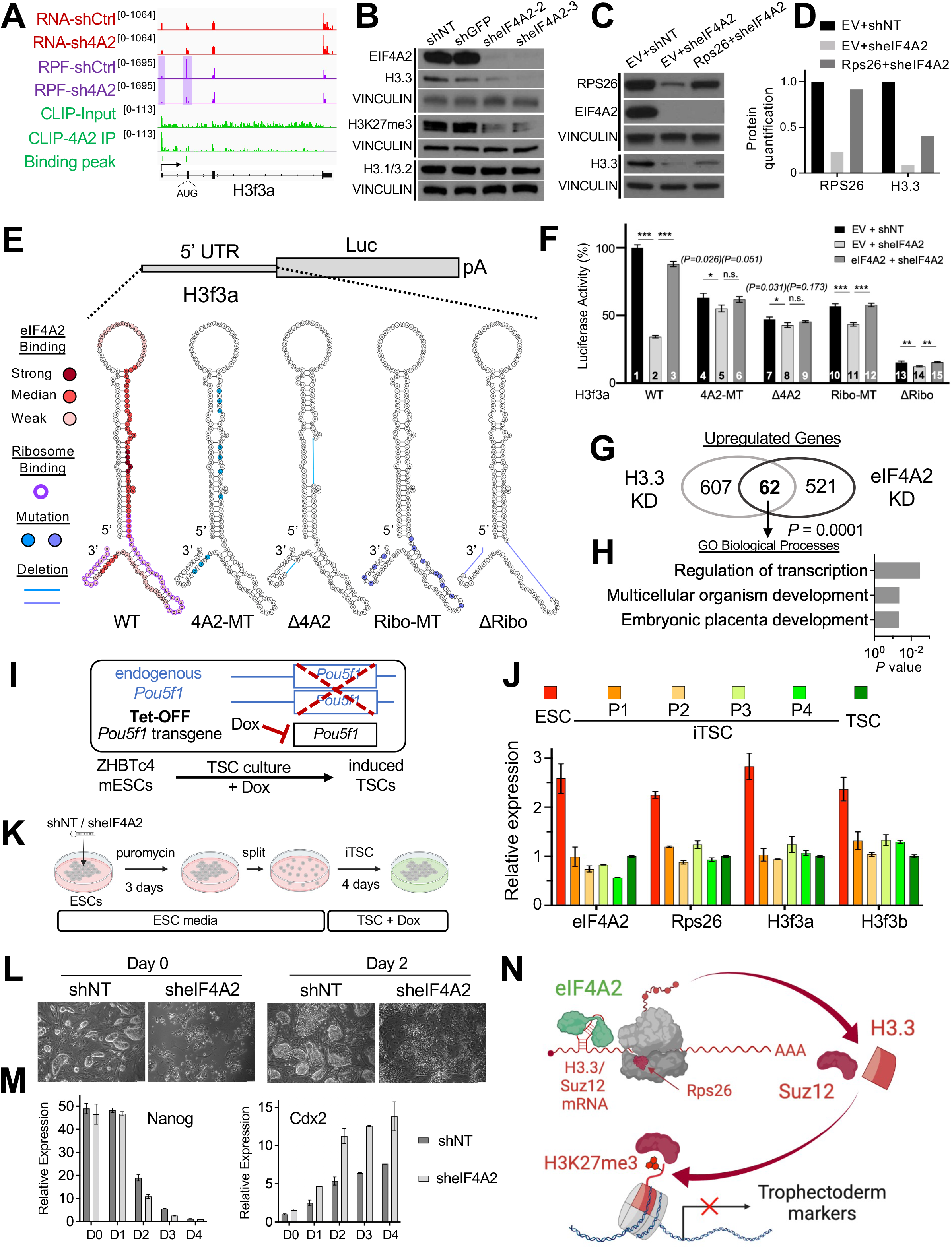
eIF4A2 activates the translation initiation of H3.3 to inhibit trophectoderm differentiation. (**A**) IGV snapshots on H3f3a with similar datasets are shown in Fig. 2H. The arrow indicates the transcription start direction. The start codon is located in the second exon. (**B**) Western blots of the indicated proteins between control and eIF4A2 KD ESCs. VINCULINs serve as a loading control. (**C** and **D**) Western blots of the indicated proteins in ESCs expressing the indicated shRNA and plasmid (C). The protein quantification (D) is normalized to the VINCULIN protein density. (**E**) (Top) Luciferase reporter constructs in which the luciferase gene is driven by H3f3a 5’ UTR WT or mutants. (Bottom) The secondary structure of the H3f3a 5’ UTR WT or mutants. WT: wildtype. MT: the mutant with mutations that disrupt the eIF4A2 (4A2-MT) or ribosome (Ribo-MT) binding region. Δ4A2/ΔRibo: the mutant with the deletion of eIF4A2 (Δ4A2) or ribosome (ΔRibo) binding site. In WT, red indicates eIF4A2 binding with the gradient denoting the binding strength; purple indicates the ribosome binding. In mutants, blue/purple indicates the mutation or the deletion in the eIF4A2 (blue)/ribosome (purple) binding region. (**F**) Luciferase activity of mRNAs driven by the H3f3a 5’ UTR WT or mutants in ESCs with the indicated shRNA and plasmid. (**G** and **H**) Venn diagrams showing upregulated transcripts upon eIF4A2 KD or H3.3 KD in ESCs (G) with corresponding GO terms of overlapped genes (H). (**I**) Illustration of TSC induction from ZHBTc4 ESCs by doxycycline treatment (Tet-OFF transgenic *Pou5f1* encoding Oct4 protein) and medium change to TSC culture. (**J**) qRT-PCR of indicated transcripts during the induction of TSCs from ZHBTc4 ESCs, relative to the expression levels in TSC, normalized to β-actin expression. P1-4: passage number, three days per passage. TSC: embryo-derived trophoblast stem cells. (**K**) Schematic of TSC induction from ZHBTc4 ESCs with control KD (shNT) or eIF4A2 KD (sheIF4A2). (**L**) Cell morphology of iTSC induction at Day 0 and Day 2, with shNT or sheIF4A2. (**M**) qRT-PCR of Nanog and Cdx2 during the induction of TSCs from ZHBTc4 ESCs, with shNT or sheIF4A2. (**N**) A model depicting eIF4A2-mediated translation initiation activation of H3.3 in the inhibition of extraembryonic trophectoderm in ESCs.

To understand how eIF4A2 binding to H3.3 mRNAs leads to translation activation, we used multiple RNA secondary structure prediction tools (see details in Methods), which yielded similar predicted structures for H3f3a/b 5’ UTRs, revealing that most of the eIF4A2-binding sites map to structured stem-loops (Fig. 5E, and fig. S6E, and fig. S7, A and B). Next, we performed the luciferase reporter assays with either mutations (MT) or deletions (Δ) of the eIF4A2 binding regions or ribosome binding regions in H3f3a/b 5’ UTRs (Fig. 5, E and F, and fig. S6, E and F, and fig. S7). Our results (Fig. 5F and fig. S6F) revealed: first, all mutants caused various reductions in luciferase activity relative to WT [compare 4 (the bar number, same below), 7, 10, 13 with 1], indicating that both eIF4A2- and ribosome-binding sites are critical for efficient translation; second, upon eIF4A2 KD, a much larger reduction of luciferase activity was observed for WT reporter (compare 1 with 2) than any other individual mutants (compare each black bar with each light gray bar, such as 4 with 5), indicating that eIF4A2 loss only minimally exacerbates the translational defect already present in each individual mutants; third, upon eIF4A2 KD, ectopic expression of shRNA-immune eIF4A2 cDNA can rescue the translational defects caused by the loss of eIF4A2 expression in the WT reporters (compare 2 with 3) or the reporters with the mutations/deletions of ribosomal binding sites (compare 11 with 12, 14 with 15), but not of eIF4A2 binding regions (compare 5 with 6, 8 with 9). These results are consistent with a general requirement of ribosome binding in translation, and more importantly, highlight a specific contribution of eIF4A2 and its 5’ UTR binding to the efficient translation of H3.3.

Depletion of H3.3 was reported to de-repress trophoblast lineage genes in ESCs (fig. S6G) through disengagement of PRC2/SUZ12 (another Rps26-dependent target, Fig. 4, C and D) and consequent downregulation of H3K27me3 (*8*), which was also observed in eIF4A2-depleted ESCs (Fig. 1H) with the decreased H3K27me3 level (Fig. 5B). Comparison of upregulated genes in H3.3 KD (*8*) and eIF4A2 KD revealed a significant number of overlapped genes with enrichment of GO terms such as “embryonic placenta development” (Fig. 5, G and H), consistent with the GSEA results (Fig. 1H and fig. S6G). Furthermore, downregulation of eIF4A2, H3.3, and Rps26 was observed during the induction of trophoblast stem cells (iTSCs) from Oct4-depleted ESCs (*6*) (Fig. 5, I and J, and fig. S6, H and I). The depletion of eIF4A2 also facilitated the differentiation of ESCs to iTSCs (Fig. 5K) based on the changes of both the morphology (Fig. 5L) and expression of the pluripotency and TSC markers (Fig. 5M). Finally, analysis of the in vivo expression data (*34*) also revealed higher expression levels of eIF4A2, H3f3a, and H3f3b in the blastocyst than the placenta (Rps26 is not available) (fig. S6J), supportive to our in vitro findings.

These results indicate that eIF4A2 participates in Rps26-dependent ribosomes to activate the translation initiation of a specific histone variant H3.3 and together with Suz12, restricting trophectoderm differentiation and safeguarding ESC identity (Fig. 5N).

### eIF4A2 activates and interacts with Ddx6 to inhibit the totipotency 2-cell marker Zscan4 RNA and protein

Apart from roles in translation activation, eIF4A2 is also responsible for translation repression of many differentiation-promoting genes such as Hexim1 (*25*), Tcf12 (*26*), Tcf3 (*27*), and Tet1 (*28*) (Fig. 3A bottom). Given that Zscan4 was the most upregulated protein (Fig. 2F) with a concomitant increase of the double-positive (Zscan4+/MERVL+) 2C-reporter (*35*) populations in eIF4A2 KD ESCs (Fig. 6, A to C) and its prominent roles in regulating totipotent 2-cell like cells (*5, 36*), we decided to address how eIF4A2 represses Zscan4 in ESCs.

**Fig. 6:**
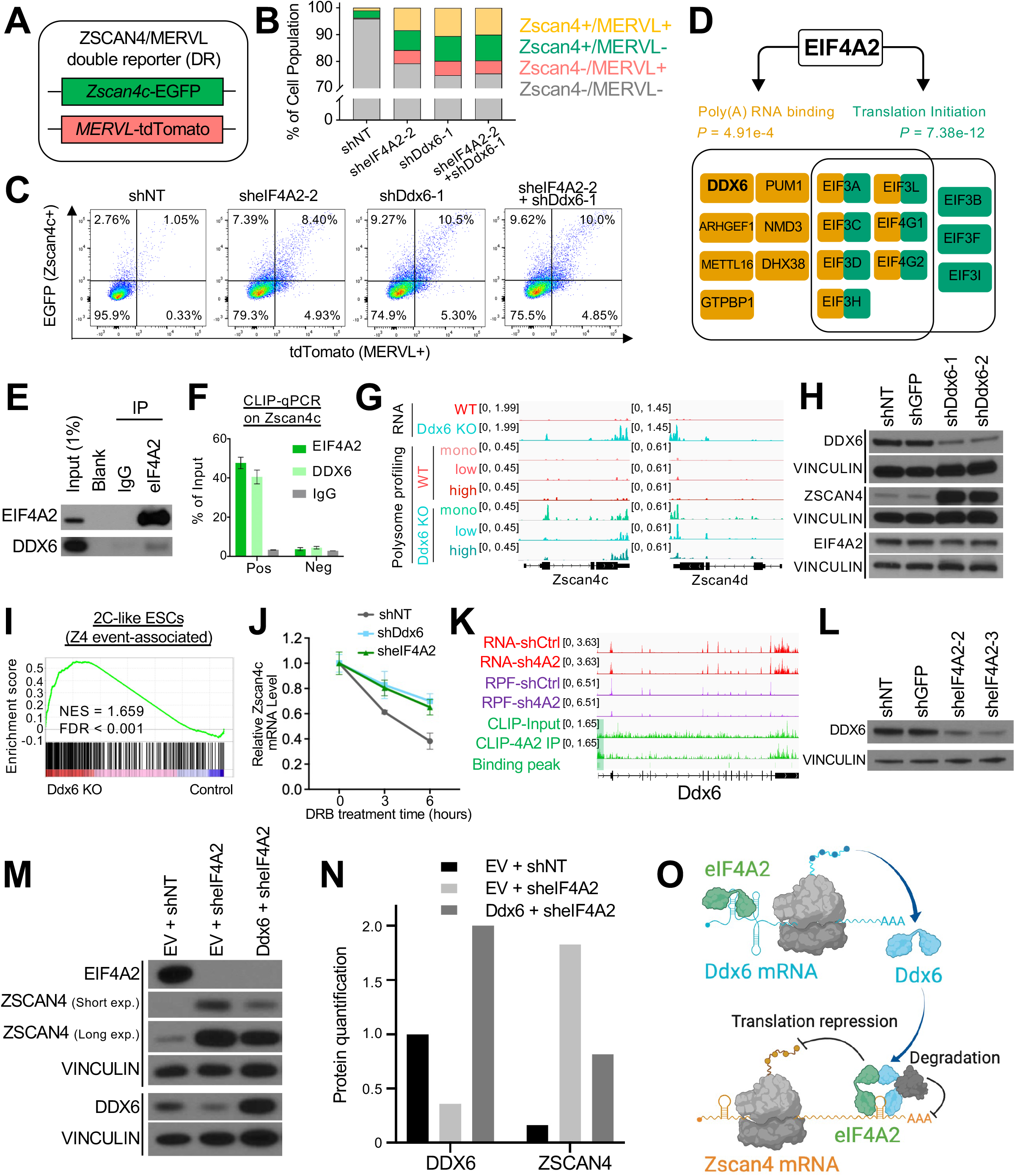
eIF4A2 represses the 2-cell gene Zscan4 through the interaction with Ddx6. (**A**) Depiction of the double-reporter system with *Zscan4c-*EGFP and *MERVL-*tdTomato. (**B** and **C**) The flow cytometry (B) and the quantification (C) results of indicated populations (Zscan4+/MERVL+, Zscan4+/MERVL-, Zscan4-/MERVL+, Zscan4-/MERVL-) among control KD (shNT), eIF4A2 KD, Ddx6 KD, and double (eIF4A2 and Ddx6) KD. (**D**) The top two GO terms of the EIF4A2 interactome in ESCs, indicated with the involved proteins. (**E, H**, and **L**) Western blots of the indicated proteins under treatments as indicated. VINCULIN serves as the loading control (H and L). (**F**) CLIP-qPCR on Zscan4c with EIF4A2, DDX6, or IgG pulldown. The amplicon positions are labeled in Fig. 2H. (**G**) IGV snapshots on Zscan4c/d showing RNA-seq data (top two), polysome profiling data for monosome (mono), low polysome (low), and high polysome (high) (middle three and bottom three) between control WT (middle three) and *Ddx6* KO (bottom three). RNA-seq and polysome profiling data are from GSE112765 and GSE112761 (*38*). (**I**) GSEA result of the 2C-like ESCs (Z4 event: Zscan4 expression) gene set by comparing *Ddx6* KO with control cells. (**J**) qRT-PCR of Zscan4c in shNT-, shDdx6-, or sheIF4A2-infected ESCs with the treatment of the transcription inhibitor DRB (5,6-Dichlorobenzimidazole 1-β-D- ribofuranoside, 100 μM) at different time points. (**K**) IGV snapshots on Ddx6 with similar datasets are shown in Fig. 2H. (**M** and **N**) Western blots of the indicated proteins in ESCs expressing the indicated shRNA and plasmid (M). The protein quantification (N) is normalized to the Vinculin protein density. (**O**) A model depicting eIF4A2-mediated translation initiation activation of Ddx6 and the cooperation between eIF4A2 and Ddx6 in the inhibition of Zscan4 in ESCs.

Zscan4 is a well-established marker of the 2C totipotency program (*36*). To confirm that the double-positive (Zscan4+/MERVL+) cell subpopulation upon eIF4A2 KD were real 2-cell like cells but not an intermediate cell population co-expressing the 2C and differentiation markers, we sorted the double-positive (Zscan4+/MERVL+) and double-negative (Zscan4-/MERVL-) ESC subpopulations upon eIF4A2 KD or control KD (two replicates for each group) and performed the RNA-seq (fig. S8A). The PCA result showed that the double-positive groups upon eIF4A2 KD were the closest to the double-positive groups upon control KD (shNT) (fig. S8B). And the GSEA result revealed that the double-positive groups upon eIF4A2 KD even displayed a stronger 2C-like ESC gene expression signature compared with the double-positive groups upon control KD (fig. S8C). Examples of 2C marker genes (Zscan4c, Zscan4d, and USP7la; fig. S8, D and G) also confirmed that eIF4A2 KD enhanced the 2C-like ESC gene expression signature in the double-positive subpopulation, supporting the repressive roles of eIF4A2 in the control of Zscan4 expression. Furthermore, comparison between the transcriptomes of eIF4A2 KD double-positive cells and control KD double-negative cells (equivalent to the predominant pluripotent cell population in wildtype ESCs) further highlighted the enrichment of 2C gene signature but not differentiation gene signature (data not shown). These results together disfavor the possibility of Zscan4+ cell populations upon eIF4A2 KD being an intermediate cell population co-expressing 2C and differentiation genes.

The translational repression roles of eIF4A2 was explored in HeLa cells, in which EIF4A2 interacts with the components of the CCR4-NOT deadenylase complex to inhibit the translation of its targets (*37*). To examine whether a similar repressive mechanism exists in ESCs, we performed EIF4A2 immunoprecipitation followed by mass spectrometry (IP-MS) in ESCs (table S5 and fig. S8E). While many translation initiation factors were identified in the EIF4A2 interactome, supporting its roles in translation initiation, we did not detect CCR4-NOT components, suggesting a potentially different mechanism of eIF4A2 for translational repression in ESCs. Instead of CCR4-NOT members, DDX6 was identified in the EIF4A2 interactome in ESCs (Fig. 6, D and E), which was reported to play roles in mRNA degradation and translational repression in ESCs (*38, 39*). We confirmed the co-binding of DDX6 and EIF4A2 on Zscan4c mRNA (Fig. 2H and Fig. 6F). Both RNA and protein upregulation of Zscan4c/d were observed from RNA-seq and polysome profiling of Ddx6-depleted ESCs compared with WT ESCs (*38*) (Fig. 6G). Ddx6 KD increased ZSCAN4 protein level (with a minimal effect on EIF4A2 protein level) (Fig. 6H), the ZSCAN4-positive cell population (fig. S8F), and double-positive populations in *Zscan4c*-EGFP/*MERVL*-tdTomato double reporter ESCs (Fig. 6, A to C). Ddx6 depletion also upregulated Zscan4 expression (i.e., Z4 event (*17*))-associated 2C-like signature transcripts (Fig. 6I). These data support the involvement of Ddx6 in eIF4A2-mediated Zscan4 repression.

Zscan4 expression is highly heterogeneous in ESC culture (*36*). To examine expression profiling of eIF4A2, Ddx6, and Zscan4 in ESCs, we first analyzed their RNA expression levels in mouse early embryogenesis, revealing that both eIF4A2 and Ddx6 have relatively low expression levels during the 2C stage (fig. S1D and fig. S8G). In ESC culture, the single-cell RNA-sequencing (scRNA-seq) (*40*) indicates that, compared with the expression patterns of Oct4 (relatively homogeneous), Nanog (relatively heterogeneous/mosaic-in-colony), and Zscan4c (highly heterogeneous/spot-in-colony) (*41*), both eIF4A2 and Ddx6 are expressed in a rather heterogeneous pattern like Nanog (fig. S8H). However, when we ranked the cells based on Zscan4c RNA expression level (UMIFM counts from 0 to 4), neither eIF4A2 nor Ddx6 showed obvious Zscan4c correlated or anti-correlated RNA expression pattern (fig. S8I), suggesting the important post-transcriptional (including translational) control of these factors. We then performed immunofluorescence to examine their protein levels, revealing relatively heterogeneous expression patterns of EIF4A2 and DDX6 and confirming the P-body location of DDX6 in ESCs as reported (*38, 39*) (fig. S8J). Zscan4+ cells are not present in the cell population with P-bodies (indicated by Ddx6-expressing dots in fig. S8J, middle panels). However, we did notice a small number of non-P-body DDX6-expressing cells co-express ZSCAN4 with the expression of DDX6 in a diffused instead of classical P-body dotted pattern (Zoom-in bottom panels in fig. S8J). We further confirmed that Ddx6 depletion inhibited the degradation of Zscan4c mRNA (Fig. 6J). These results suggest a specific role of Ddx6 in the repression of Zscan4 via a P-body-dependent manner (and possibly also a P-body independent role of Ddx6 in Zscan4 activation).

To lend further support that eIF4A2 and Ddx6 may act together on gene repression, we compared upregulated genes in eIF4A2 KD (Fig. 1F) and *Ddx6* KO cells (*38*). We observed a significant number of co-repressed genes enriched in “cell differentiation” and “multicellular organism development” (fig. S8, K and L), consistent with the differentiation phenotype of both eIF4A2 KD (Fig. 1 and fig. S1) and Ddx6 KD ESCs (*38*) (fig. S8, M and N). However, unlike eIF4A2 KD (Fig. 2, H and I), Ddx6 KD didn’t downregulate the protein levels of OCT4/NANOG/SOX2 (fig. S8, O and P). Interestingly, EIF4A2 binds to the TIR of Ddx6 mRNA and translationally activates Ddx6, supported by the maintained mRNA abundance but decreased RPF levels of Ddx6 upon eIF4A2 KD (Fig. 6K). Based on the RPF change pattern, Ddx6 is an Rps26-independent target under eIF4A2-mediated translation initiation activation, confirmed by the luciferase reporter assays and western blot of DDX6 in eIF4A2- or Rps26-depleted ESCs (Fig. 4, C, G, and H, and Fig. 6L). And eIF4A2 KD also prevented degradation of Zscan4c mRNA, consistent with Ddx6 depletion (Fig. 6J). Moreover, while double KD of eIF4A2 and Ddx6 didn’t synergize the upregulation of the 2C-like population compared with the single depletion (Fig. 6, A-C), overexpression of Ddx6 partially but not fully rescued the upregulation of ZSCAN4 protein level in eIF4A2-depleted ESCs (Fig. 6, M and N), suggesting that eIF4A2 and Ddx6 may function through the same pathway to regulate the 2C-like subpopulation in ESCs, and that additional factors other than Ddx6 may also contribute to eIF4A2-mediated repression of Zscan4 abundance (see Discussion).

Together, these results establish a model whereby eIF4A2 restricts the totipotency 2C program by activating the translation initiation of Ddx6 mRNA and recruiting DDX6 protein for Zscan4 mRNA degradation and translational repression in maintaining pluripotency of ESCs (Fig. 6O).

## Discussion

We demonstrate that eIF4A2 is specifically responsible for a unique translation initiation control mechanism dedicated to safeguarding ESC identity through restricting embryonic/extraembryonic differentiation and the totipotency 2C program (Fig. 7). On the one hand, EIF4A2 activates the translation initiation of histone variant H3.3 (together with Polycomb protein Suz12) and pluripotency factors possibly through Rps26-dependent and -independent mechanisms, respectively, distinguished by RNA characteristics or regulon (i.e., RNA sequence elements and structural complexity) surrounding the start codon and the 5’ UTR, leading to the activation of pluripotency program and repression of multilineage differentiation (Fig. 7A). On the other hand, EIF4A2 translationally activates Ddx6 mRNA and interacts with DDX6 in binding CDS near 3’ UTR of Zscan4 mRNAs, leading to Zscan4 mRNA degradation and translation repression in restricting totipotency 2C program in pluripotent cells (Fig. 7B). We thus established a translational paradigm in the cytoplasm for the protein synthesis of stem cell potency transcription factors (e.g., OSN, Zscan4) and epigenetic regulators (e.g., H3.3 and Suz12) imposed on their well-established transcriptional roles in the nucleus in promoting pluripotency and repressing trophectoderm/totipotency. Lending further support of our findings, eIF4A2 constitutive knockout mice were embryonic lethal (https://www.taconic.com/knockout-mouse/eif4a2-trapped) (unpublished dataset from Taconic Biosciences Inc), and both mouse (fig. S1, J and K) and human (*42*) naïve ESCs rely on eIF4A2, but not eIF4A1, dependent pathways to form a more compact naïve proteome for translating selective mRNAs (*42*).

**Fig. 7:**
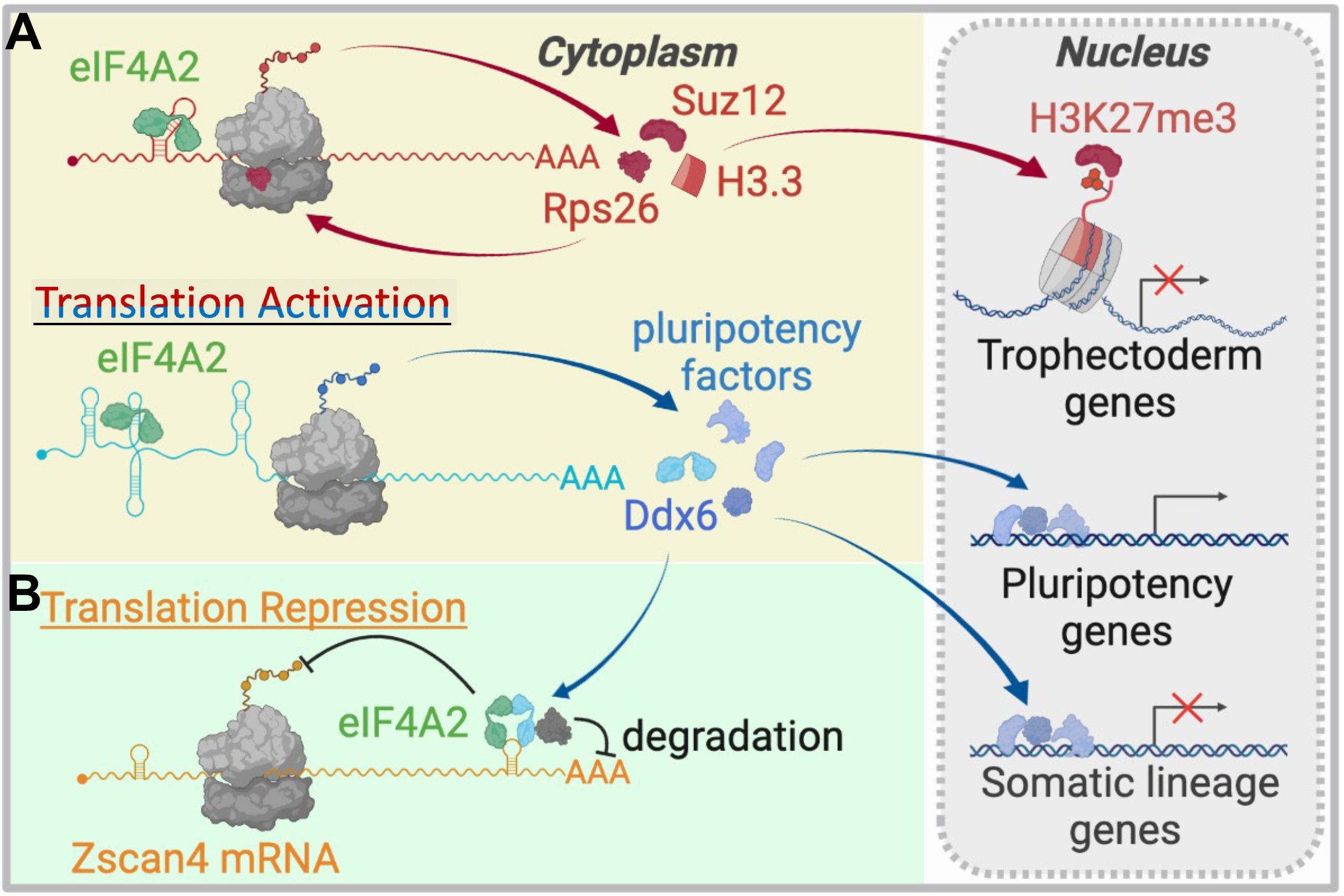
A model depicting eIF4A2-mediated translational control in safeguarding ESC identity. eIF4A2 is responsible for a unique translation initiation control network dedicated to safeguarding ESC identity. (**A**) eIF4A2 binds to the TIR of its targets to activate the translation initiation: eIF4A2 activates the translation initiation of H3.3 and Rps26 through Rps26-containing ribosomes (red); eIF4A2 also activates specific pluripotency-associated mRNAs and Ddx6 through Rps26-deficient ribosomes (blue). (**B**) Via the physical interaction with Ddx6, eIF4A2 represses Zscan4’s expression by binding CDS near 3’ UTR (non-TIR) of Zscan4 mRNAs (orange).

An outstanding question remains as to why ribosome binding is only lost at TIR instead of FL mRNA bodies upon eIF4A2 depletion for the Rps26-dependent targets. Emerging evidence shows that ribosomes can be stalled on transcripts during translation elongation (*43, 44*). mRNA translation is an energy-consuming process (*45*). 5’ TOP transcripts are highly abundant and they are best-translated by the heavy polysomes, and a small perturbation in these 5’ TOP RNA-coding protein levels can lead to an acute and profound impact on broad downstream translation events (*46, 47*). Thus, ribosome stalling can avoid wasting dedicated energy and provide a rapid response to deter the current ongoing 5’ TOP translation products. Upon eIF4A2 KD and Rps26 downregulation, Rps26-independent ribosomes of high abundance, which may be enriched in monosomes and/or light polysomes in the normal condition, may start to bind and translate the

Rps26-dependent targets, causing stalling and trigger *cis*-acting feedback inhibition of translation initiation (*48*). This can help explain the loss of ribosome binding at TIRs but no other regions on the Rps26-dependent targets upon eIF4A2 depletion. Moreover, for the differences around the start codons among the Rps26-dependent/independent targets (Fig. 3H), apart from the differences around the “-4” position, where Rps26 binds (*33*), differences were also observed at -3, -5, and +4 positions, which could be due to the overall ribosome structural changes upon Rps26 depletion altering the whole ribosome binding profile on mRNAs. Although Rps26 was reported substoichiometric in free ribosome subunits relative to heavy polysomes in mouse ESCs (*49*), it remains to be determined whether it is substoichiometric in monosomes/light polysomes and whether other ribosome proteins other than Rps26 in the eIF4A2 translatome may act alone or together with Rps26 in contributing to the functional specialization of eIF4A2 in translational control of stem cell pluripotency.

The Rps26-dependent targets also include H3.3-coding mRNAs. We didn’t detect H3.3 in the SILAC-MS assay (table S2), which may be due to the highly similar peptide sequences between H3.3 and H3.1/H3.2, as well as the highly K/R-rich histone peptides that may be over-digested by trypsin in SILAC-MS. Unlike canonical H3.1/H3.2 mRNAs with stem-loop structures at 3’ UTRs for SLBP-mediated translation (*50*), H3.3 mRNAs are non-canonical with introns, polyadenylated tails, and longer 5’ UTRs/3’ UTRs without stem-loop structures at 3’ UTRs. Such unique features potentially endow H3.3 with distinct gene expression control. Our study identified an eIF4A2-mediated translation initiation activation of H3.3 in ESCs with Rps26-dependent ribosomes. The role of H3.3 during early development and the recurrent mutations of H3.3 in multiple types of cancers have been well recognized (*51*), highlighting the importance of our findings on translation control of H3.3 in further understanding its roles in development and disease.

We showed that eIF4A2 activates the translation initiation of pluripotency-associated mRNAs through Rps26-independent ribosomes. Mettl3 and Mettl14, two components of the methyltransferase complex, are the targets underlying such translation initiation control (table S2), which deposit m^6^A on key pluripotency transcripts to promote their degradation (*52*). This raises an intriguing possibility: eIF4A2 KD decreases protein synthesis of key pluripotency factors, but their mRNA levels, which are supposed to be downregulated as well based on the feedforward transcriptional circuitry (*23*), are maintained. This can be explained by the stabilization of these mRNAs with reduced m^6^A modifications (*52*), resulting from reduced protein levels of METTL3/14 whose mRNAs are subject to eIF4A2-mediated translation initiation activation. Future investigation is needed to validate this potential mechanism.

Embryonic development is associated with the stepwise restriction of cell potency from totipotent 2C embryos to pluripotent inner cell mass of the blastocyst (*53*); accordingly, pluripotency may require a proper restriction of the totipotency 2C program in ESCs. In ESCs, the expression of Zscan4 is transient and reversible in only 1-5% of the cell population (*17*), and overexpression of Zscan4c in ESCs activates MT2/MERVL and 2C genes (including Zscan4 cluster) (*22*), indicating an elaborate mechanism in ESCs responsible for the repression of Zscan4 and the associated 2C program. Here, we presented a novel mechanism of eIF4A2-mediated repression of Zscan4 in ESCs, partly through Ddx6. The derepression of Zscan4 upon eIF4A2/Ddx6 KD emerged in only a subpopulation of ESCs (fig. S3H and fig. S8F), which may explain the low magnitude in bulk RNA expression changes of 2C markers (table S1). DDX6 can also be recruited to 5’ UTRs of some eIF4A2 TIR targets (data not shown), which warrants future investigation for their potential collaborative or competitive roles in target gene expression control. Interestingly, H3.3 also impedes the 2C program in ESCs (*54*). It remains to be determined how eIF4A2-mediated translation repression and activation of Zscan4c/d and H3.3, respectively, may crosstalk or synergize in restricting the 2C program in ESCs and early development. Finally, apart from DDX6, EIF4A2 may enlist other factors in the EIF4A2 interactome, such as PUM1, to restrict 2C and lineage differentiation in maintaining the ESC identity. Pum1 is a repressor found in P-bodies and stress granules, and it can promote ESC differentiation during exit from pluripotency (*55*). In undifferentiated ESCs, EIF4A2 may sequester PUM1 through their physical association and restrict its functions to prevent ESC differentiation, which warrants future investigation.

## Acknowledgments

We thank A.K. Hadjantonakis for sharing TSC lines, G. Yeo and E.V. Nostrand for eCLIP-seq technical advice, the ISMMS genomics core facility, NYU Langone’s genome technology center, and Illumina Tech Support for technical assistance, Jill Gregory for the illustrations (Fig. 1A and Fig. 2, B and G), a shared instrumentation grant for the LSR II Flow Cytometer (S10RR027050) to the Columbia Center for Translational Immunology (CCTI) Flow Cytometry Core, and members of the Wang lab for discussions. The research in Wang laboratory was supported by NIH (GM129157, HD095938, and HD097268) and NYSTEM (C32583GG and C32569GG).

## Author contributions

DL designed, conducted the experiments, and wrote the manuscript. DL, JY, and XH performed the data analysis. XH and HZ provided reagents and experimental support. JW conceived, designed, supervised the studies, and wrote and approved the final manuscript.

## Competing interests

The authors declare no competing interests.

## Methods

### Murine Embryonic Stem Cell Culture

Feeder-free murine ESCs were grown on 0.1% gelatin-coated plates in ESC medium containing high-glucose DMEM, 15% fetal bovine serum (FBS), 100 μM nonessential amino acids (NEAA), 1% nucleoside mix, 2 mM L-Glutamine, 50 U/mL Penicillin/Streptomycin, 0.1 mM 2-mercaptoethanol, and homemade recombinant leukemia inhibitory factor (LIF) tested for efficient self-renewal maintenance, at 37 °C and 5% CO_2_. For naïve culture conditions (2i/LIF), murine ESCs were cultured on 0.1% gelatin-coated plates using serum-free N2B27 medium (DMEM/F12 and Neurobasal media were used in a ratio of 1:1, 1x B27 supplement, 1x N2 supplement, 50 U/mL Penicillin/Streptomycin, 2 mM L-glutamine, and 0.1 mM 2-mercaptoethanol) supplemented with Gsk3β inhibitor (CHIR99021, 3 μM final) and Mek inhibitor (PD0325901, 1 μM final) and recombinant LIF.

### TSC culture and iTSC reprogramming

ZHBTc4 ESCs are cultured in serum + LIF ESC medium. For iTSC, medium was replaced to TSC medium (RPMI 1640 supplemented with 20% FBS, 50 U/mL Penicillin/Streptomycin, 0.1 mM 2-mercaptoethanol, 2 mM L-Glutamine, 100 μM nonessential amino acids, 25 ng/ml recombinant FGF4, and 1 μg/ml heparin) supplemented with 1 μg/ml doxycycline.

### Transfection and Lentiviral Infection

Transfection of cells was performed using Lipofectamine 3000 according to the manufacturer’s manual. The production of lentivirus and viral infection were performed as described (*56*). All shRNA knockdown experiments followed the same time points as the RNAi screen, which ended on Day 4.5 with 2.5-day drug selection (1 μg/mL puromycin).

### RNAi Screen

To perform a TIF RNAi screen, we selected the constitutive shRNAs (three independent shRNAs per gene) with validated KD efficiencies wherever data were available in the literature. For those that have not been reported, we selected three independent shRNAs with the best-predicted KD efficiency targeting both exons and 3’ UTR regions. The lentivirus was prepared as previously described (*56*). ESCs were seeded in the gelatin-coated tissue culture plates together with the viruses for viral infection (details as previously described) (*56*) (Day 0). On Day 1, the virus/medium was changed with regular serum/LIF ESC medium. From Day 2, the medium was changed with serum/LIF ESC medium containing 1 μg/mL puromycin daily to select the infected cells. The negative control for drug selection showed that the drug treatment killed all uninfected cells after 2 days. On Day 4.5, AP staining was performed to record the phenotype.

This screen identified eIF4A2 as the TIF specifically required for ESC maintenance from an RNAi screen of all factors involved in translation initiation. To our knowledge, this is the first screen specifically focused on the translation initiation machinery. Many large-scale RNAi/CRISPR screens collected results 48 or 72 hours after introductions of short RNAs (shRNA/siRNA/sgRNA) against targets (*57-60*), but as median half-lives of eukaryotic mRNAs and proteins are 9 hours and 46 hours separately (*61*), a lot of important candidates were overlooked due to the presence of their undegraded proteins at the end of those screens. Such timepoints are particularly important for the factors involved in translational control as it would take longer for their targets to have responses in their protein levels. Therefore, we ended our screen at day 4.5 post-infection. This timepoint is indeed important as upon eIF4A2 KD, the loss of the dome-shaped ESC morphology started around 3.5 days later, explaining why eIF4A2 was overlooked by many previous screens. In addition, we recorded screen results with AP staining, a direct phenotypic marker of pluripotent stem cells, instead of transgenic reporters used in many other studies (*57-59*). Transgenic reporters are proper and sensitive to transcriptional change while not suitable for identifying direct translational change, and they are limited to the response of certain transcriptional regulatory elements.

### iPSC Reprogramming

MEF reprogramming was performed as described (*16*) with a few modifications. Briefly, 100,000 reprogrammable MEFs containing a Dox-inducible OKSM cassette were infected with shRNA lentiviruses or transfected with plasmids, followed with 1 μg/mL puromycin or 250 μg/mL Hygromycin. After the drug selection, 95,000 MEFs were seeded on top of irradiated MEF feeders on a 12-well tissue culture plate coated with gelatin, in Dox-containing serum/LIF ESC medium (day 0). From day 13, the medium was changed to serum/LIF ESC medium without Dox until day 19, when AP-staining was performed to record the reprogramming result.

MEF-derived Nanog-GFP pre-iPSCs were maintained and used for reprogramming as described (*62*). Briefly, MEF-derived Nanog-GFP pre-iPSCs were infected with shRNA lentiviruses or transfected with plasmids. 20,000 pre-iPSCs were seeded after selection on a 12-well tissue culture plate coated with gelatin and grown in serum + LIF for 2 days before medium switch to 2i + LIF. On day 10 in 2i + LIF, Nanog-GFP iPSC colonies were counted under fluorescence microscopy.

### Immunofluorescence Staining

Cells were fixed with 4% paraformaldehyde (w/v) for 15 min at room temperature (RT), washed, then permeabilized with 0.25% Triton X-100 solution for 5 min at RT and blocked with 5% fetal bovine serum (FBS). Then cells were incubated with primary antibodies and 5% FBS in PBS overnight at 4 °C. The next day, after washing, cells were incubated with secondary antibodies and 3 μM DAPI with 5% FBS in PBS for 1 hour at RT in the dark. After washing, cells were imaged with a Leica DMI 6000 inverted microscope.

### Whole-Cell Extract Preparation, Co-Immunoprecipitation, and Western Blot

Whole-cell lysates were harvested from ESCs in Lysis Buffer [50 mM HEPES (pH 7.6), 250 mM NaCl, 0.1% NP-40, 0.2 mM EDTA, 1.4 mM β-mercaptoethanol, 0.2 mM PMSF, 1x protease inhibitor cocktail] and before IP, NaCl concentration of the lysates was diluted to 179 mM with Dilution Buffer [20 mM Tris (pH 7.6), 20% glycerol, 0.05% NP-40, 0.2 mM EDTA, 1.4 mM β-mercaptoethanol, 0.2 mM PMSF, 1x protease inhibitor cocktail]. The lysates were incubated with the anti-eIF4A2 antibody or IgG control by rotating overnight at 4°C. On the second day, Protein G agarose beads were equilibrated with Lysis Buffer diluted with Dilution Buffer (179 mM NaCl). The lysate/antibody mixtures were added to the equilibrated beads and rotated for 3 hours at 4°C. The bound beads were washed five times with Wash Buffer [50 mM HEPES (pH 7.6), 179 mM NaCl, 0.1% NP-40, 0.2 mM EDTA, 1.4 mM β-mercaptoethanol, 0.2 mM PMSF] and then eluted with SDS-DTT loading buffer by boiling for 5 min at 95 °C. Eluted proteins were separated by SDS-PAGE and visualized by western blotting. True-blot secondary antibody was used to reduce the IgG detection.

### Immunoprecipitation of EIF4A2 Protein Complexes in ESCs and LC-MS/MS Analysis

To identify EIF4A2 interacting partners in ESCs, we used three different anti-EIF4A2 antibodies [IP1: ab194471 (Mouse polyclonal, Abcam), IP2: ab31218 (Rabbit polyclonal, Abcam), IP3: PA5-27431 (Rabbit polyclonal, ThermoFisher)] to isolate EIF4A2 protein complexes independently for mass spectrometry (MS) identification, with IgG pull-down as controls (for ab194471, the control was a normal mouse IgG polyclonal antibody, 12-371 from Sigma-Aldrich; for ab31218 and PA5-27431, the control was a normal rabbit IgG polyclonal antibody, PP64 from Millipore Sigma). Whole-cell lysates were prepared as previously described from 20×15 cm dishes of WT ESCs. Then to decrease the salt concentration to 100 mM, the lysates were transferred to a dialyzer with Dialysis Buffer [20 mM HEPES (pH 7.9), 20% glycerol (v/v), 100 mM KCl, 1.5 mM MgCl_2_, 0.2 mM EDTA, 0.5 mM DTT, 0.2 mM PMSF, 1x protease inhibitor cocktail] at 4°C for 3 hours. The precipitated proteins were removed by centrifugation. Then the total protein mass of the lysates was determined after protein concentration measurement. 10 mg proteins were used for each IP, and some lysates were left as Input. IP lysates were diluted with IP-DNP buffer (Dialysis Buffer + 0.02% NP-40) to 12 mL, and Benzonase (Pierce, 20 μL 15 U/μL) was added to remove DNA and RNA. The lysates were pre-cleared with 100 μL IP-DNP buffer-equilibrated Protein G agarose beads per 10 mg total protein for 1 hour at 4°C, followed by incubation with 20 μg anti-eIF4A2 antibody (or IgG) by rotating overnight at 4°C. On the second day, Protein G agarose beads were equilibrated with IP-DNP buffer. The lysate/antibody mixtures were added to the equilibrated beads and rotated for 3 hours at 4°C. After the bound beads were washed five times with IP-DNP buffer, the protein complexes were eluted with SDS loading buffer by boiling for 5 min at 95°C and separated by an SDS-PAGE gel. Whole lanes were excised and subjected to liquid chromatography-tandem mass spectrometry (LC-MS/MS) analysis. MS data were processed by Thermo Proteome Discoverer software with SEQUEST engine against Swiss-Prot mouse protein sequence database. Proteins were filtered by the minimal number of identified unique peptides (>=2). Common contamination proteins (e.g., keratins) were removed, and spectral count (PSM, the number of peptide spectrum matches) ratio of (EIF4A2 IP/IgG) >=4 was applied. The list was cleaned with CRAPome (*63*). The lists for three IP/IgG groups with spectral count ratios are present in table S5. Out of the three lists, the proteins presented in at least two lists were combined as the final EIF4A2 interactome in ESCs for GO analysis (Fig. 6D).

### SILAC-MS Profiling of Relative Protein Levels

ESCs were cultured in the medium labeled by either light (L-arginine and L-lysine) or heavy (L-^13^C_6_^15^N_4_-arginine and L-^13^C_6_^15^N_2_-lysine) for more than two weeks. The cells cultured in light and heavy medium were infected with control shRNAs and eIF4A2 shRNAs, respectively. The infected cells were selected by puromycin (1 μg/mL), and the cell lysates at different SILAC mediums were equally mixed as indicated in Fig. 2E, resulting in 4 mixtures. Protein lysates were dissolved in 8M Urea buffer and subjected to tryptic digestion, followed by LC-MS/MS using an Orbitrap-Velos mass spectrometer. MS data were processed by Thermo Proteome Discoverer software for protein quantification and identification. Proteins were filtered by being identified in at two out of four replicates (“Count” >= 2). All the data and statistical results, including the standard deviation (SD) and relative standard deviation (RSD), are present in table S2.

### Apoptosis Detection Assay and Flow Cytometry

Apoptotic analysis was determined using the FITC Annexin V apoptosis detection kit with PI (Propidium Iodide) (Biolegend 640914) and performed according to the manufacturer’s manual. First, single-cell suspension was achieved by using cell strainers to remove large clumps of cells. Then both Annexin V and PI-stained cells were analyzed by flow cytometry. The flow cytometry used an LSR-II Flow Cytometer (BD Biosciences) and data were analyzed using FlowJo software.

### Protein Synthesis Measurements Using OP-Puromycin Incorporation

To measure protein synthesis, ESCs were incubated for 30 min in serum/LIF medium supplemented with O-propargyl-puromycin (OP-Puro, 50 μM, Abcam, ab146664). Cells were then harvested, washed with PBS, fixed with 4% paraformaldehyde for 15 minutes on ice, and permeabilized with PBS supplemented with 3% FBS for 5 minutes at room temperature. The Click-iT Plus OP-Puro Alexa Fluor 647 assay was done according to the manufacturer’s protocol (Click-iT Cell Reaction Buffer Kit, ThermoFisher Scientific, C10269). Cells were resuspended in 200 μL PBS supplemented with 3% FBS and 0.1% saponin and analyzed by flow cytometry. To inhibit OP-Puro incorporation (the CHX group), cycloheximide (CHX, 100 μg/mL) was added 30 min before OP-Puro.

### Luciferase Assay

Luciferase assay was performed in ESCs transfected with 10 ng pRL-TK and 200 ng luciferase reporter plasmids containing the 5’ UTR elements using Lipofectamine 3000. Forty-eight hours after transfection, the cells were lysed, and luminescence was assayed using the Dual-Glo luciferase assay kit (Promega, E2920) according to the manufacturer’s manual. The measurements were performed in triplicate biological samples.

### RNA Extraction and qRT-PCR

RNA was extracted from the indicated cell lines with the RNeasy kit (Qiagen, 74136) and converted to cDNA using qScript (Quanta, 95048). Relative gene expression levels were analyzed with Lightcycler 480 SYBR green master mix (Roche, 4729749001) on the LightCycler480 real-time PCR system (Roche). Gene expression levels were normalized to the β-actin expression level.

### RNA-seq and Data Analysis

RNA-seq was performed in ESCs infected with control shRNAs or eIF4A2 shRNAs. Biological duplicates were prepared. Total RNA from each sample was extracted from the cells with the RNeasy kit (Qiagen, 74136). Samples were prepared, indexed, pooled, and sequenced on the Illumina HiSeq system according to a PolyA selection protocol per the manufacturer’s instructions.

RNA-seq reads were aligned to the mouse mm9 genome using Bowtie2 (v2.3.4.3), and aligned bam files were sorted by name using the parameter -n. We used the HTSeq software (v0.11.2) and mm9 annotation file from GENCODE (https://www.gencodegenes.org/mouse/release_M1.html) to count reads for each gene using parameters -r name -f bam, and BioMart (*64*) to retrieve corresponding genes names. Finally, Read counts were normalized with the trimmed mean of M-values (TMM) method (*65*) for differential expression analysis using edgeR (v3.26.8) (*66*).

Public RNA-seq data were downloaded (refer to Data availability), aligned to mm9, and then followed with the same processing setting as mentioned above.

The significance value in Fig. 5G and fig. S8K reflects the probability of finding overlapping genes using the hypergeometric test.

### RNA Secondary Structure Prediction

RNA secondary structures were determined using: (1) RNAfold web server with minimum free energy prediction and thermodynamic ensemble prediction (http://rna.tbi.univie.ac.at/cgi-bin/RNAWebSuite/RNAfold.cgi); (2) RNAstructure (https://rna.urmc.rochester.edu/RNAstructureWeb/Servers/Predict1/Predict1.html) with Fold results, MaxExpect results, and ProbKnot results; (3) vs_subopt (version 5.39) (http://www.rna.it-chiba.ac.jp/~vsfold/vs_subopt/). The energy of stem-loop, stack, exterior-loop, and bulge-loop are computed by the software RNAstruture(v6.2) (https://rna.urmc.rochester.edu/RNAstructure.html). For each gene, only the structure with the lowest total energy is taken into consideration.

### GO and GSEA

Gene ontology (GO) analysis was carried out by the DAVID (The Database for Annotation, Visualization, and Integrated Discovery) functional annotation program (https://david.ncifcrf.gov/home.jsp). The terms are ranked according to the *P-value* that the program provided with default parameters.

Gene set enrichment analysis (GSEA, v4.1.0, available at https://www.gsea-msigdb.org/gsea/index.jsp) was used to determine whether the set was statistically enriched in eIF4A2 KD versus control KD, Ddx6 KO versus WT (*38*), and H3.3 KD versus control KD (*8*). The 2C-like ESC (Z4 event-associated) gene set was from a published RNA-seq dataset containing significantly more highly expressed genes in Zscan4^+^ cells than in Zscan4^-^ cells (*17*). The other gene sets were from the GSEA MSigDB database (the Molecular Signatures Database): Epidermis development (systematic name: M14065), Mesoderm development (systematic name: M15421), Endoderm differentiation (systematic name: M34153), Formation of primary germ layer (systematic name: M10670), Placenta genes (systematic name: M16071), Hindbrain differentiation (systematic name: M13307), and Cell differentiation (GO: 0030154). The normalized enrichment score (NES) and FDR were calculated by GSEA and indicated for each enrichment test.

### eCLIP-seq and Data Analysis

eCLIP-seq libraries were performed in duplicates according to the published eCLIP-seq protocol (*20*). Briefly, 20 million ESCs were UV crosslinked at 400 mJ/cm^2^ with 254 nm radiation. Cells were lysed in iCLIP lysis buffer and sonicated with Bioruptor. The cell lysates were treated with diluted RNase I to fragment RNA. The eIF4A2 antibody (Abcam, ab31218) was pre-coupled to Protein G Dynabeads and then added into the cell lysate, followed by overnight incubation at 4°C. 2% of the lysate was taken as the input sample, and the remaining lysate was magnetically separated and washed with lysis buffer. During washing, RNA was dephosphorylated with FastAP and T4 PNK, followed by a 3’RNA adapter ligation with T4 RNA ligase. Then Protein-RNA complexes were separated by an SDS-PAGE gel, transferred to nitrocellulose membranes. eCLIP was performed by excising the membrane area based on the molecular weight of eIF4A2 from the site of their molecular weight (47 kDa) to the site with 75 kDa more (122 kDa). The SMInput (size-matched input) was used for each biological replicate, and the excised area was the same as its corresponding IP sample. The details of RNA adapter ligation, immunoprecipitation, western blot, RNA purification, reverse transcription, DNA adapter ligation, cDNA quantification, PCR amplification, library purification were performed as eCLIP-seq protocol (*20*). eCLIP-seq reads were processed by following the Nature Method protocol (*20*). Adapters were trimmed (cutadapt v1.18), and reads less than 18 bp were discarded using parameters -m 18 -a NNNNNNNNNNAGATCGGAAGAGCACACGTCTGAACTCCAGTCAC -g ACGCTCTTCCGATCT -A AGATCGGAAGAGCGT -A GATCGGAAGAGCGTC -A ATCGGAAGAGCGTCG -A TCGGAAGAGCGTCGT -A CGGAAGAGCGTCGTG -A

GGAAGAGCGTCGTGT for round 1 and parameters -m 18 -A AGATCGGAAGAGCGT -A GATCGGAAGAGCGTC -A ATCGGAAGAGCGTCG -A TCGGAAGAGCGTCGT –A CGGAAGAGCGTCGTG -A GGAAGAGCGTCGTGT for round 2. Mapping reads were then performed against mouse elements in RepBase (*67*) with STAR (v2.6.1b) (*68*). Repeat mapping reads were segregated, and all others were mapped against the mouse mm9 genome with STAR (v2.6.1b). PCR duplicates were removed from uniquely mapping reads to get usable reads. Multiple in-line barcodes were merged for usable reads, followed by peak identification with the clipper software (v0.2.0, https://pypi.org/project/clipper/) using parameters -s mm9 --Bonferroni --superlocal --threshold-method binomial --save-pickle on read 2 only (Bonferroni correction was employed on our peaks to reduce false-positives. A semi-experimental option, “-superlocal”, was used to pick up peaks that may be missed with genome-wide or gene-wide thresholds). In addition, based on these candidate peaks, we further use the eCLIP size-matched input as control and then compare eCLIP with the input data to get peaks enriched in eCLIP samples with an intensity fold change of more than 2. Given these, we used the same code “-s mm9 --Bonferroni --superlocal -- threshold-method binomial --save-pickle” for peak calling and “Peak_input_normalization_wrapper.pl” for peak normalization in the above-mentioned Nature method paper, and then filtered with fold change > 2 to get our peaks. TIRs were extracted by the code (https://github.com/stephenfloor/extract-transcript-regions). Binding peaks on the coding exon and TIR were extracted as the target- and TIR target-binding peaks. These peaks were then annotated with gene names using annotatePeaks.pl script in HOMER tools (*69*), respectively. Genes in the former list were identified as targets, and those in the latter were TIR targets.

For repetitive analysis, both reads were counted for input and eCLIP data on 3’ UTR, intron, CDS, and 5’ UTR regions and then normalized by the TMM method (*65*). Fold changes were calculated for both replicates using the TMM value. The correlation coefficient was finally calculated based on fold change values to indicate the data repeatability. For peak distribution analysis, we used the Guitar (v1.20.1) (*70*) to visualize the binding frequency on 5’ UTR, CDS, and 3’ UTR regions.

### Ribosome Profiling and Data Analysis

Ribosome profiling was performed using ESCs infected with control shRNAs (shNT, shGFP) or eIF4A2 shRNAs. Total RNA and RPF libraries were prepared using the TruSeq Ribo Profile (Mammalian) Kit (Illumina, RPHMR12126) according to the manufacturer’s reference guide (document #15066016 v01) with Ribo-Zero Gold Kit (H/M/R) (Illumina, MRZG126). The prepared libraries were sequenced on a HiSeq 2500 system (Illumina).

Ribosome profiling data reads were firstly adapter trimmed using FASTX-Toolkit (v0.0.14; http://hannonlab.cshl.edu/fastx_toolkit) with the parameter -a AGATCGGAAGAGCACACGTCT. Then, rRNA and tRNA reads were removed using Bowtie (v1.2.3). The remained reads are aligned to the mouse mm9 genome using TopHat (v2.1.1). The matched total RNA-seq data were also processed with the same processing procedure. These aligned bam files were sorted by name with the parameter –n and counted by HTSeq (v0.11.2). For genes, we counted using parameters -r name -f bam. For transcripts, we used parameters -r name -f bam --nonunique all to count reads on TIR (Translation initiation region) and CDS regions. All read counts were normalized using the TMM method (*65*).

In Fig. 3A, to select the candidates through which eIF4A2 controls the translation initiation program to safeguard ESC identity, we applied the following stringent criteria to filter the eIF4A2 targets [from the middle panel (table S3) to the bottom panel (table S4) in Fig. 3A]: (1) the mRNAs are targeted by eIF4A2 at TIR; (2) upon eIF4A2 KD, RPF changes on both TIR and full-length RNA have a similar trend (both increase or both decrease); (3) the mRNAs with low ribosome density are excluded to obtain the most accurate regulated changes instead of the variance caused by noises (the targets with either of the following conditions were excluded: a, the average TMM of RPF on FL RNA in all samples <15; b, the average TMM of RPF on RNA TIR in all samples / TIR length (nt) < 0.02). The pluripotency-associated targets (fig. S3E) are not in the group of translational change (Fig. 3A, bottom) is due to their RPF foldchanges at TIRs not crossing the threshold 0.5 (higher than 0.5).

### Polysome Profiling Data Analysis

Public polysome profiling data were downloaded (GEO; Accession: GSE112761) and then trimmed using Trim Galore (v0.5.0). Trimmed data were aligned to the mm9 genome using bowtie2 (v2.3.4.3), then sorted with parameter -n and counted using HTSeq software (v0.11.2). All read counts were normalized using the TMM method.

### Sequence Enrichment Analysis

Information for all mm9 mRNA transcripts was extracted from the Ensembl database (Release 67, http://may2012.archive.ensembl.org/Mus_musculus/Info/Index), including 5’ UTR length, 5’ UTR sequence, CDS length, CDS sequence. The Minimum free energy was calculated for all 5’ UTR sequences using RNAalifold (v2.4.11) (*71*). We conducted sequence motif analysis on 5’ UTR sequences, TIR binding sequences, and ribosome binding sequences around the start codon, and these results were visualized by the ggseqlogo (v0.1) (*72*).

### Motif Enrichment Analysis

RNA motifs were determined using the findMotifsGenome.pl script in HOMER tools (*69*) with the parameter -rna.

### High-Throughput Sequencing Data Visualization

All processed and index sorted bam files of high-throughput sequencing data were converted to TDF files using the count command of igvtools, followed by visualization using IGV software (*73*).

### Quantification and Statistical Analysis

Except for GSEA, all other statistical analysis was performed with GraphPad Prism (GraphPad Software, Inc.), Excel or R (www.r-project.org/). Statistical significance was identified by Student’s *t*-test or one-way ANOVA with Tukey’s post-test as indicated in the manuscript or figure legends. *P* values of less than 0.05 were considered statistically significant. *\*P*<0.05, *\*\*P*< 0.01, \*\*\**P*< 0.001.

## Data Availability

The RNA-seq, Ribo-seq, and eCLIP-seq datasets generated during this study are available at NCBI Gene Expression Omnibus (GEO): GSE150682. Previously published data reanalyzed here are available under accession codes GSE66390, GSE76505, GSE112765, GSE112761, GSE42154.

## Notes

### Competing Interest Statement

The authors have declared no competing interest.

## References

1. T. W. Theunissen, R. Jaenisch, Mechanisms of gene regulation in human embryos and pluripotent stem cells. Development (Cambridge, England) 144, 4496–4509 (2017).

2. D. Li, M. S. Kishta, J. Wang, Regulation of pluripotency and reprogramming by RNA binding proteins. Current topics in developmental biology 138, 113–138 (2020).

3. J. A. Hackett et al., Tracing the transitions from pluripotency to germ cell fate with CRISPR screening. Nature Communications 9, (2018).

4. J. A. Saba, K. Liakath-Ali, R. Green, F. M. Watt, Translational control of stem cell function. Nat Rev Mol Cell Biol, (2021).

5. T. S. Macfarlan et al., Embryonic stem cell potency fluctuates with endogenous retrovirus activity. Nature 487, 57–63 (2012).

6. H. Niwa, J. I. Miyazaki, A. G. Smith, Quantitative expression of Oct-3/4 defines differentiation, dedifferentiation or self-renewal of ES cells. Nature Genetics 24, 372–376 (2000).

7. H. Niwa et al., Interaction between Oct3/4 and Cdx2 determines trophectoderm differentiation. Cell 123, 917–929 (2005).

8. L. A. Banaszynski et al., Hira-dependent histone H3.3 deposition facilitates PRC2 recruitment at developmental loci in ES cells. Cell 155, 107–120 (2013).

9. C. E. Aitken, J. R. Lorsch, A mechanistic overview of translation initiation in eukaryotes. Nature structural & molecular biology 19, 568–576 (2012).

10. Y. Gao et al., Protein Expression Landscape of Mouse Embryos during Pre-implantation Development. CellReports 21, 3957–3969 (2017).

11. N. Y. Chia et al., A genome-wide RNAi screen reveals determinants of human embryonic stem cell identity. Nature 468, 316–320 (2010).

12. H. Sugiyama et al., Nat1 promotes translation of specific proteins that induce differentiation of mouse embryonic stem cells. Proceedings of the National Academy of Sciences of the United States of America 114, 340–345 (2017).

13. T. Hart et al., High-Resolution CRISPR Screens Reveal Fitness Genes and Genotype-Specific Cancer Liabilities. Cell 163, 1515–1526 (2015).

14. Q. L. Ying et al., The ground state of embryonic stem cell self-renewal. Nature 453, 519–523 (2008).

15. A. Pause, N. Sonenberg, Mutational analysis of a DEAD box RNA helicase: the mammalian translation initiation factor eIF-4A. Embo J 11, 2643–2654 (1992).

16. S. E. Vidal, B. Amlani, T. Chen, A. Tsirigos, M. Stadtfeld, Combinatorial modulation of signaling pathways reveals cell-type-specific requirements for highly efficient and synchronous iPSC reprogramming. Stem Cell Reports 3, 574–584 (2014).

17. T. Akiyama et al., Transient bursts of Zscan4 expression are accompanied by the rapid derepression of heterochromatin in mouse embryonic stem cells. DNA Res 22, 307–318 (2015).

18. G. Galicia-Vázquez, R. Cencic, F. Robert, A. Q. Agenor, J. Pelletier, A cellular response linking eIF4AI activity to eIF4AII transcription. RNA (New York, N.Y.) 18, 1373–1384 (2012).

19. G. Galicia-Vázquez, J. Chu, J. Pelletier, eIF4AII is dispensable for miRNA-mediated gene silencing. RNA (New York, N.Y.) 21, 1826–1833 (2015).

20. E. L. Van Nostrand et al., Robust transcriptome-wide discovery of RNA-binding protein binding sites with enhanced CLIP (eCLIP). Nature Methods 13, 508–514 (2016).

21. K. Leppek, R. Das, M. Barna, Functional 5’ UTR mRNA structures in eukaryotic translation regulation and how to find them. Nat Rev Mol Cell Biol 19, 158–174 (2018).

22. W. Zhang et al., Zscan4c activates endogenous retrovirus MERVL and cleavage embryo genes. Nucleic Acids Research 47, 8485–8501 (2019).

23. R. A. Young, Control of the embryonic stem cell state. Cell 144, 940–954 (2011).

24. M. Giacomello, A. Pyakurel, C. Glytsou, L. Scorrano, The cell biology of mitochondrial membrane dynamics. Nat Rev Mol Cell Biol 21, 204–224 (2020).

25. V. Ding et al., HEXIM1 induces differentiation of human pluripotent stem cells. PloS one 8, e72823 (2013).

26. S. Mesman, M. P. Smidt, Tcf12 Is Involved in Early Cell-Fate Determination and Subset Specification of Midbrain Dopamine Neurons. Front Mol Neurosci 10, 353 (2017).

27. J. Wray et al., Inhibition of glycogen synthase kinase-3 alleviates Tcf3 repression of the pluripotency network and increases embryonic stem cell resistance to differentiation. Nat Cell Biol 13, 838–845 (2011).

28. L. Cimmino, O. Abdel-Wahab, R. L. Levine, I. Aifantis, TET family proteins and their role in stem cell differentiation and transformation. Cell Stem Cell 9, 193–204 (2011).

29. O. Meyuhas, T. Kahan, The race to decipher the top secrets of TOP mRNAs. Biochimica et Biophysica Acta (BBA) - Gene Regulatory Mechanisms 1849, 801–811 (2015).

30. R. Yamashita, S. Sugano, Y. Suzuki, K. Nakai, DBTSS: DataBase of Transcriptional Start Sites progress report in 2012.

31. C. C. Thoreen et al., A unifying model for mTORC1-mediated regulation of mRNA translation. Nature 485, 109–113 (2012).

32. D. Li, J. Wang, Ribosome heterogeneity in stem cells and development. The Journal of cell biology 219, (2020).

33. M. B. Ferretti, H. Ghalei, E. A. Ward, E. L. Potts, K. Karbstein, Rps26 directs mRNA-specific translation by recognition of Kozak sequence elements. Nature Structural & Molecular Biology 24, 700–707 (2017).

34. A. P. Hutchins et al., Models of global gene expression define major domains of cell type and tissue identity. Nucleic Acids Res 45, 2354–2367 (2017).

35. F. Yang et al., DUX-miR-344-ZMYM2-Mediated Activation of MERVL LTRs Induces a Totipotent 2C-like State. Cell Stem Cell 26, 234–250 e237 (2020).

36. G. Falco et al., Zscan4: a novel gene expressed exclusively in late 2-cell embryos and embryonic stem cells. Dev Biol 307, 539–550 (2007).

37. A. Wilczynska et al., eIF4A2 drives repression of translation at initiation by Ccr4-Not through purine-rich motifs in the 5’UTR. Genome Biol 20, 262 (2019).

38. J. W. Freimer, T. J. Hu, R. Blelloch, Decoupling the impact of microRNAs on translational repression versus RNA degradation in embryonic stem cells. eLife 7, (2018).

39. B. Di Stefano et al., The RNA Helicase DDX6 Controls Cellular Plasticity by Modulating P-Body Homeostasis. Cell Stem Cell 25, 622-638.e613 (2019).

40. A. M. Klein et al., Droplet barcoding for single-cell transcriptomics applied to embryonic stem cells. Cell 161, 1187–1201 (2015).

41. M. G. Carter et al., An in situ hybridization-based screen for heterogeneously expressed genes in mouse ES cells. Gene Expr Patterns 8, 181–198 (2008).

42. C. Chen et al., Translational and Post-translational Control of Human Naïve versus Primed Pluripotency. iScience, 103645 (2021).

43. S. Das Sharma et al., Widespread Alterations in Translation Elongation in the Brain of Juvenile Fmr1 Knockout Mice. Cell Reports 26, 3313-3322.e3315 (2019).

44. E. W. Mills, R. Green, Ribosomopathies: There’s strength in numbers. Science 358, (2017).

45. N. Lane, W. Martin, The energetics of genome complexity. Nature 467, 929–934 (2010).

46. A. Bulut-Karslioglu et al., The Transcriptionally Permissive Chromatin State of Embryonic Stem Cells Is Acutely Tuned to Translational Output. Cell Stem Cell 22, 369-383.e368 (2018).

47. X. Pichon et al., RNA binding protein/RNA element interactions and the control of translation. Curr Protein Pept Sci 13, 294–304 (2012).

48. S. Juszkiewicz et al., Ribosome collisions trigger cis-acting feedback inhibition of translation initiation. Elife 9, (2020).

49. Z. Shi et al., Heterogeneous Ribosomes Preferentially Translate Distinct Subpools of mRNAs Genome-wide. Molecular Cell 67, 71-83.e77 (2017).

50. W. F. Marzluff, E. J. Wagner, R. J. Duronio, Metabolism and regulation of canonical histone mRNAs: life without a poly(A) tail NIH Public Access. Nat Rev Genet 9, 843–854 (2008).

51. L. Shi, H. Wen, X. Shi, The Histone Variant H3.3 in Transcriptional Regulation and Human Disease. J Mol Biol 429, 1934–1945 (2017).

52. F. Aguilo et al., Coordination of m(6)A mRNA Methylation and Gene Transcription by ZFP217 Regulates Pluripotency and Reprogramming. Cell Stem Cell 17, 689–704 (2015).

53. C. Chazaud, Y. Yamanaka, Lineage specification in the mouse preimplantation embryo. Development 143, 1063–1074 (2016).

54. Q. Tian, X. F. Wang, S. M. Xie, Y. Yin, L. Q. Zhou, H3.3 impedes zygotic transcriptional program activated by Dux. Biochemical and biophysical research communications 522, 422–427 (2020).

55. A. C. Goldstrohm, T. M. T. Hall, K. M. McKenney, Post-transcriptional Regulatory Functions of Mammalian Pumilio Proteins. Trends Genet 34, 972–990 (2018).

56. N. Ivanova et al., Dissecting self-renewal in stem cells with RNA interference. Nature 442, 533–538 (2006).

57. J. Betschinger et al., Exit from pluripotency is gated by intracellular redistribution of the bHLH transcription factor Tfe3. Cell 153, 335–347 (2013).

58. T. G. Fazzio, J. T. Huff, B. Panning, An RNAi Screen of Chromatin Proteins Identifies Tip60-p400 as a Regulator of Embryonic Stem Cell Identity. Cell 134, 162–174 (2008).

59. C. Schaniel et al., Smarcc1/Baf155 Couples Self-Renewal Gene Repression with Changes in Chromatin Structure in Mouse Embryonic Stem Cells. Stem Cells 27, N/A-N/A (2009).

60. L. Ding et al., Systems Analyses Reveal Shared and Diverse Attributes of Oct4 Regulation in Pluripotent Cells. Cell Systems 1, 141–151 (2015).

61. B. Schwanhäusser et al., Global quantification of mammalian gene expression control. Nature 473, 337–342 (2011).

62. A. Saunders et al., The SIN3A/HDAC Corepressor Complex Functionally Cooperates with NANOG to Promote Pluripotency. Cell reports 18, 1713–1726 (2017).

63. D. Mellacheruvu et al., The CRAPome: A contaminant repository for affinity purification-mass spectrometry data. Nature Methods 10, 730–736 (2013).

64. S. Durinck et al., BioMart and Bioconductor: a powerful link between biological databases and microarray data analysis. Bioinformatics 21, 3439–3440 (2005).

65. M. D. Robinson, A. Oshlack, A scaling normalization method for differential expression analysis of RNA-seq data. Genome Biology 11, R25–R25 (2010).

66. D. J. McCarthy, Y. Chen, G. K. Smyth, Differential expression analysis of multifactor RNA-Seq experiments with respect to biological variation. Nucleic Acids Res 40, 4288–4297 (2012).

67. W. Bao, K. K. Kojima, O. Kohany, Repbase Update, a database of repetitive elements in eukaryotic genomes. Mob DNA 6, 11 (2015).

68. A. Dobin et al., STAR: ultrafast universal RNA-seq aligner. Bioinformatics 29, 15–21 (2013).

69. S. Heinz et al., Simple combinations of lineage-determining transcription factors prime cis-regulatory elements required for macrophage and B cell identities. Mol Cell 38, 576–589 (2010).

70. X. Cui et al., Guitar: An R/Bioconductor Package for Gene Annotation Guided Transcriptomic Analysis of RNA-Related Genomic Features. Biomed Res Int 2016, 8367534 (2016).

71. S. H. Bernhart, I. L. Hofacker, S. Will, A. R. Gruber, P. F. Stadler, RNAalifold: improved consensus structure prediction for RNA alignments. BMC Bioinformatics 9, 474 (2008).

72. O. Wagih, ggseqlogo: a versatile R package for drawing sequence logos. Bioinformatics 33, 3645–3647 (2017).

73. J. T. Robinson et al., Integrative genomics viewer. Nat Biotechnol 29, 24–26 (2011).

