## Supplemental Text and Figures for "eIF4A2 targets developmental potency and histone H3.3 transcripts for translational control of stem cell pluripotency"

**This PDF file includes:**

Legends for Tables S1 to S6
Figs. S1 to S7

**Other Supplementary Materials for this manuscript include the following:**

Tables S1 to S6

**Supplementary Materials**

**Tables S1-S6**

**Table S1.** RNA-Seq expression data in eIF4A2 KD and control KD ESCs. Related to Fig. 1.

**Table S2.** eIF4A2 target mRNAs identified by eCLIP-Seq in ESCs (including all the targets and the TIR targets) and SILAC data comparing protein expression levels between eIF4A2 KD and control KD. Related to Fig. 2.

**Table S3.** Ribosome profiling data with RNA, RPF, TIR_RPF levels in eIF4A2 KD and control KD ESCs. Related to Figs. 2 and 3.

**Table S4.** Transcriptome and translation initiation "regulome" of eIF4A2's TIR targets. Related to Fig. 3.

**Table S5.** eIF4A2 interactors identified by IP-Mass Spectrometry in ESCs. Related to Fig. 6D and fig. S7A.

**Table S6.** Lists of shRNAs and primers used in the study.

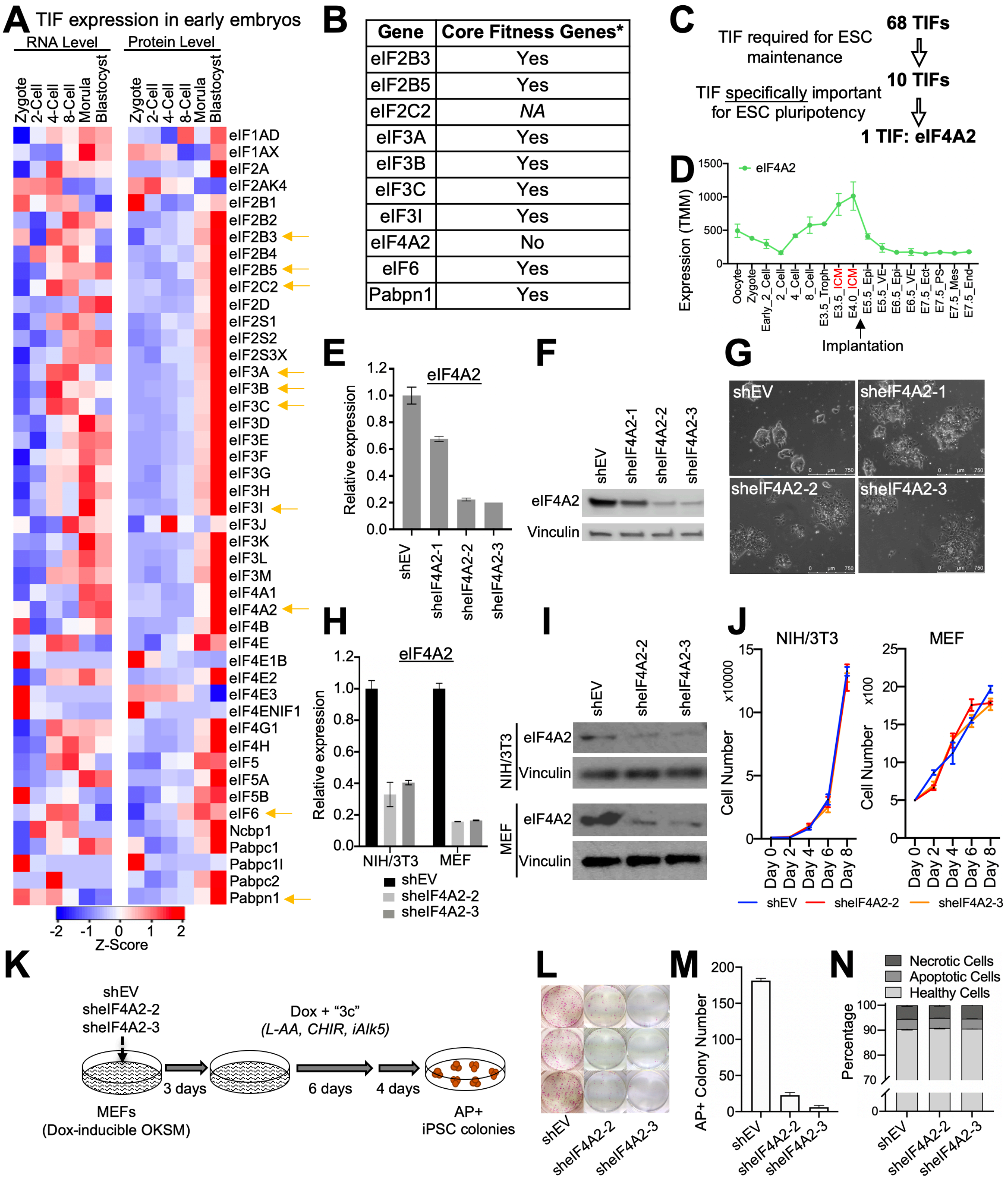
A TIF expression in early embryos
RNA Level
Protein Level
Zygote
2-Cell
4-Cell
8-Cell
Morula
Blastocyst
eIF1AD
eIF1AX
eIF2A
eIF2AK4
eIF2B1
eIF2B2
eIF2B3
eIF2B4
eIF2B5
eIF2C2
eIF2D
eIF2S1
eIF2S2
eIF2S3X
eIF3A
eIF3B
eIF3C
eIF3D
eIF3E
eIF3F
eIF3G
eIF3H
eIF3I
eIF3J
eIF3K
eIF3L
eIF3M
eIF4A1
eIF4A2
eIF4B
eIF4E
eIF4E1B
eIF4E2
eIF4E3
eIF4ENIF1
eIF4G1
eIF4H
eIF5
eIF5A
eIF5B
eIF6
Ncbp1
Pabpc1
Pabpc1l
Pabpc2
Pabpn1
-2 -1 0 1 2
Z-Score
B
Gene | Core Fitness Genes*
eIF2B3 | Yes
eIF2B5 | Yes
eIF2C2 | NA
eIF3A | Yes
eIF3B | Yes
eIF3C | Yes
eIF3I | Yes
eIF4A2 | No
eIF6 | Yes
Pabpn1 | Yes
C
TIF required for ESC maintenance
68 TIFs
TIF specifically important for ESC pluripotency
10 TIFs
1 TIF: eIF4A2
D
eIF4A2
Expression (TMM)
1500
1000
500
0
Oocyte
Zygote
Early_2_Cell
2_Cell
4_Cell
8_Cell
E3.5_Troph
E3.5_ICM
E4.0_ICM
E5.5_Epi
E5.5_VE
E6.5_Epi
E6.5_VE
E7.5_Ect
E7.5_Mes
E7.5_PS
E7.5_End
Implantation
E
eIF4A2
Relative expression
1.2
1.0
0.8
0.6
0.4
0.2
0.0
shEV
sheIF4A2-1
sheIF4A2-2
sheIF4A2-3
F
shEV
sheIF4A2-1
sheIF4A2-2
sheIF4A2-3
eIF4A2
Vinculin
G
shEV
sheIF4A2-1
sheIF4A2-2
sheIF4A2-3
H
eIF4A2
Relative expression
1.2
1.0
0.8
0.6
0.4
0.2
0.0
NIH/3T3
MEF
shEV
sheIF4A2-2
sheIF4A2-3
I
shEV
sheIF4A2-2
sheIF4A2-3
NIH/3T3
eIF4A2
Vinculin
MEF
eIF4A2
Vinculin
J
NIH/3T3
x10000
Cell Number
15
10
5
0
MEF
x100
Cell Number
25
20
15
10
5
0
Day 0
Day 2
Day 4
Day 6
Day 8
shEV
sheIF4A2-2
sheIF4A2-3
K
shEV
sheIF4A2-2
sheIF4A2-3
MEFs
(Dox-inducible OKSM)
3 days
Dox + "3c"
(L-AA, CHIR, iAlk5)
6 days
4 days
AP+
iPSC colonies
L
shEV
sheIF4A2-2
sheIF4A2-3
M
AP+ Colony Number
200
150
100
50
0
shEV
sheIF4A2-2
sheIF4A2-3
N
Necrotic Cells
Apoptotic Cells
Healthy Cells
Percentage
100
90
80
70
0
shEV
sheIF4A2-2
sheIF4A2-3

**Fig. S1. TIFs have changing expression levels during early embryo development and eIF4A2 is required for both establishment and maintenance of ESCs.**

**A**, TIF mRNA and protein expression levels in early embryos. The yellow arrows point to the expression level change of the selected candidates from the RNAi screen. **B**, Core fitness classification of candidates (*core fitness classification). **C**, The summary of the RNAi screen result. **D**, The RNA expression of eIF4A2 during mouse embryogenesis. Data is from GSE76505 and GSE66390. **E**,**F**, qPCR (E) and western blot (F) analyses to validate eIF4A2 KD efficiency with control shRNA (shEV) or eIF4A2 shRNAs. Vinculin is the loading control in western blot (F). **G**, Cell morphology of CCE ESCs with either control shRNA (shEV) or three eIF4A2 shRNAs. **H**,**I**, qRT-PCR (H) and western blot (I) analyses of eIF4A2 in NIH/3T3 and MEF cell lines with control KD and eIF4A2 KD. All qRT-PCR in Extended Data Fig. 1 is relative to the expression levels in shEV, normalized to β-actin expression. Vinculin is the loading control in western blot (I). **J**, Proliferation curves for NIH/3T3 and MEF cell lines with control KD or eIF4A2 KD. **K**, Schematic of the doxycycline (Dox)-inducible MEF reprogramming system. **L**,**M**, Representative whole-well images (L) and the quantification (M) of AP+ iPSC colonies. **N**, Apoptosis analysis of ESCs with control KD and eIF4A2 KD.

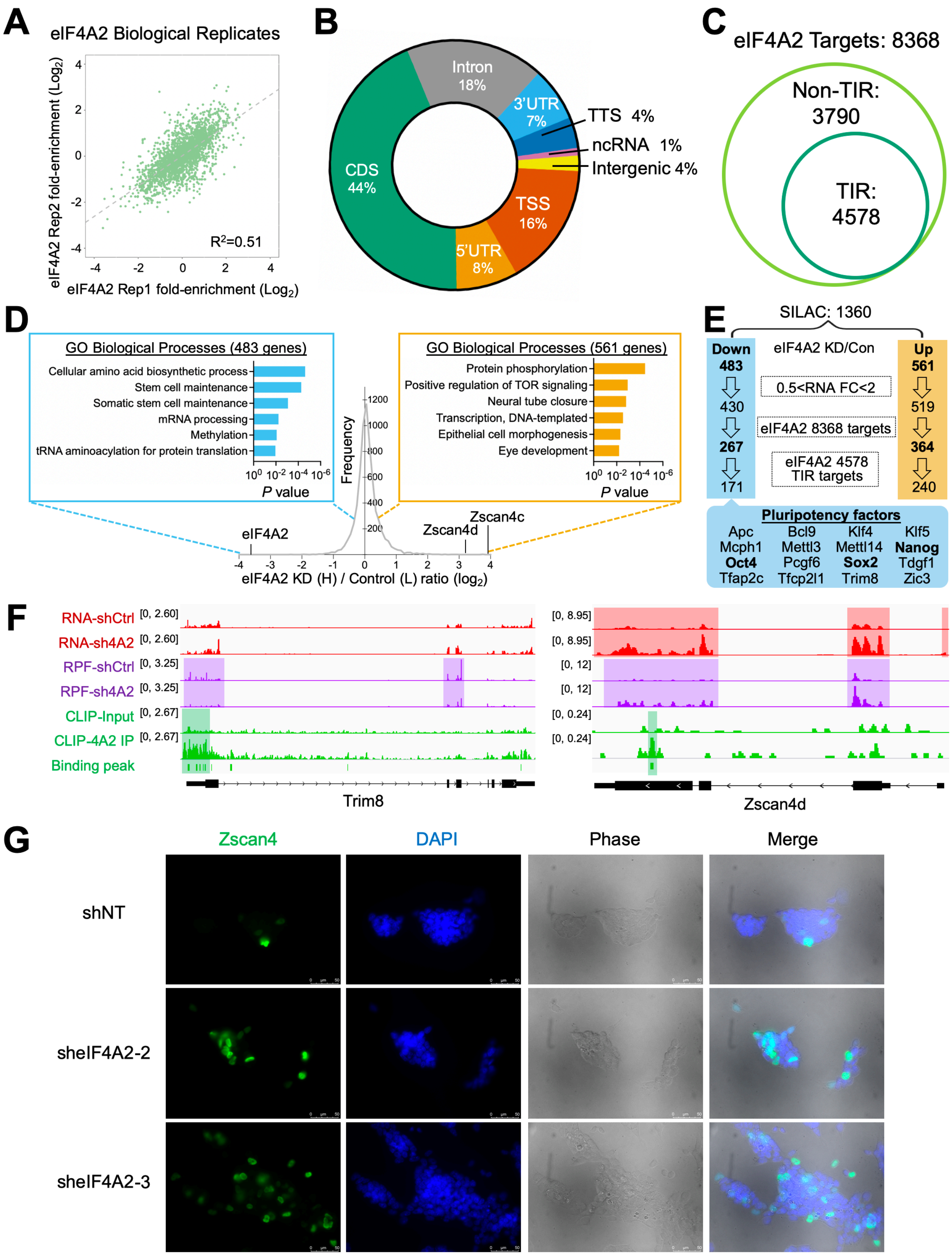
A
eIF4A2 Biological Replicates
eIF4A2 Rep2 fold-enrichment (Log2)
eIF4A2 Rep1 fold-enrichment (Log2)
R2=0.51
B
Intron 18%
3'UTR 7%
TTS 4%
ncRNA 1%
Intergenic 4%
TSS 16%
5'UTR 8%
CDS 44%
C
eIF4A2 Targets: 8368
Non-TIR: 3790
TIR: 4578
D
GO Biological Processes (483 genes)
Cellular amino acid biosynthetic process
Stem cell maintenance
Somatic stem cell maintenance
mRNA processing
Methylation
tRNA aminoacylation for protein translation
P value
GO Biological Processes (561 genes)
Protein phosphorylation
Positive regulation of TOR signaling
Neural tube closure
Transcription, DNA-templated
Epithelial cell morphogenesis
Eye development
P value
Frequency
eIF4A2
Zscan4d
Zscan4c
eIF4A2 KD (H) / Control (L) ratio (log2)
E
SILAC: 1360
Down 483
eIF4A2 KD/Con
Up 561
430
0.5<RNA FC<2
519
267
eIF4A2 8368 targets
364
171
eIF4A2 4578 TIR targets
240
Pluripotency factors
Apc Bcl9 Klf4 Klf5
Mcph1 Mettl3 Mettl14 Nanog
Oct4 Pcgf6 Sox2 Tdgf1
Tfap2c Tfcp2l1 Trim8 Zic3
F
RNA-shCtrl [0, 2.60]
RNA-sh4A2 [0, 2.60]
RPF-shCtrl [0, 3.25]
RPF-sh4A2 [0, 3.25]
CLIP-Input [0, 2.67]
CLIP-4A2 IP [0, 2.67]
Binding peak
Trim8
[0, 8.95]
[0, 8.95]
[0, 12]
[0, 12]
[0, 0.24]
[0, 0.24]
Zscan4d
G
Zscan4
DAPI
Phase
Merge
shNT
sheIF4A2-2
sheIF4A2-3

**Fig. S2. eIF4A2 is a positive and negative translation regulator that controls mRNAs encoding cellular potency factors.**

**A**, Replication analysis of eIF4A2 eCLIP-seq with two biological replicates. **B**, Pie chart indicates the distribution of significantly enriched eIF4A2 peak locations from eCLIP-seq. Please note that, in Fig. 2C, the size of 5'UTR, CDS, and 3'UTR represents their true length in the transcriptome (5'UTR is much shorter than CDS and 3'UTR), so, Fig. 2C shows a higher enrichment of peaks at 5'UTR than 3'UTR while their peak number percentages are similar here. **C**, Venn diagrams showing eIF4A2 targets and TIR targets identified by eIF4A2 eCLIP-seq. **D**, Graph of the frequency distribution of heavy/light (H/L, eIF4A2 KD/control) ratios of all proteins identified by SILAC-MS, with GO analysis of the genes with decreased (left side, blue box) or increased (right side, orange box) protein levels upon eIF4A2 KD. **E**, Flow chart showing the filtering process to identify the genes underlying eIF4A2-mediated control. **F**, IGV snapshots on the indicated genes with similar datasets are shown in Fig. 2H. **G**, Immunofluorescence of endogenous Zscan4 in ESCs with control shRNA (shNT) or eIF4A2 shRNAs.

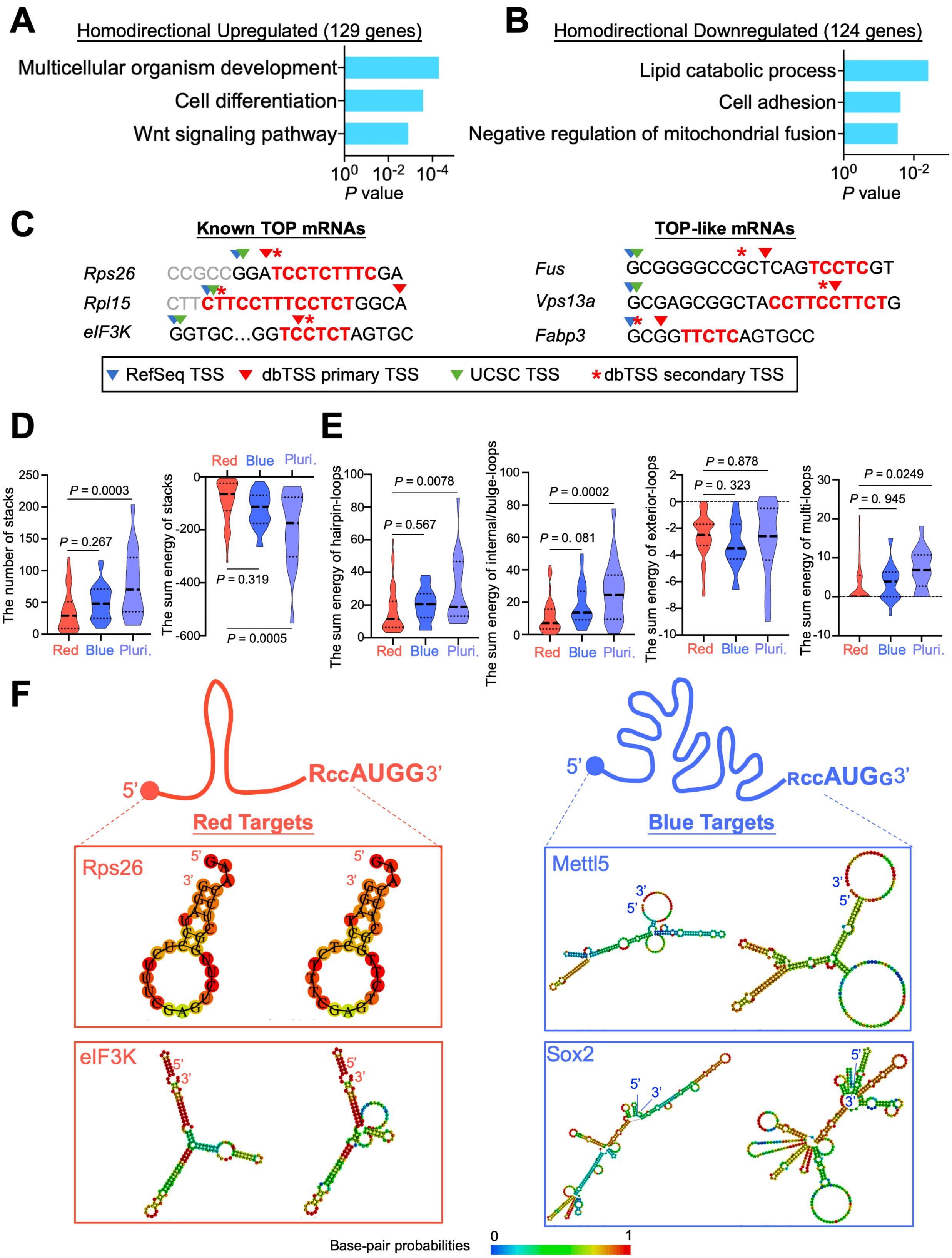
A
Homodirectional Upregulated (129 genes)
Multicellular organism development
Cell differentiation
Wnt signaling pathway
P value
B
Homodirectional Downregulated (124 genes)
Lipid catabolic process
Cell adhesion
Negative regulation of mitochondrial fusion
P value
C
Known TOP mRNAs
Rps26 CCGCCGGATCCTCTTTCGA
Rpl15 CTTCTTCCTTTCCTCTGGCA
eIF3K GGTGC…GGTCCTCTAGTGC
TOP-like mRNAs
Fus GCGGGGCCGCTCAGTCCTCGT
Vps13a GCGAGCGGCTACCTTCCTTCTG
Fabp3 GCGGTTCTCAGTGCC
RefSeq TSS
dbTSS primary TSS
UCSC TSS
dbTSS secondary TSS
D
The number of stacks
P = 0.0003
P = 0.267
The sum energy of stacks
P = 0.319
P = 0.0005
Red Blue Pluri.
E
The sum energy of hairpin-loops
P = 0.0078
P = 0.567
The sum energy of internal/bulge-loops
P = 0.0002
P = 0. 081
The sum energy of exterior-loops
P = 0.878
P = 0. 323
The sum energy of multi-loops
P = 0.0249
P = 0. 945
F
RccAUGG3'
Red Targets
Rps26
eIF3K
RccAUGG3'
Blue Targets
Mettl5
Sox2
Base-pair probabilities
0
1

**Fig. S3. eIF4A2 activates translation initiation in two modes of action depending on the RNA elements and 5'UTR complexity.**

**A**,**B**, GO analyses of homodirectional upregulated (A) or downregulated (B) genes in Fig. 3A top. **C**, TSS (transcription start site) annotations for selected TOP and TOP-like mRNAs. Primary and secondary TSS locations from dbTSS are indicated with red triangles and asterisks. TSS locations from RefSeq and UCSC are indicated with blue and green triangles. The C/T stretches are highlighted in red. **D**, The number and the sum energy of RNA stacks in 5'UTR of the three indicated target groups. **E**, The sum energy of different types of RNA loops in 5'UTR of the three indicated target groups. The results in D,E are computed by the software RNAstructure (details in Methods). **F**, The red target mRNAs contain shorter 5' UTR with simple secondary structures and stringent Kozak sequence around the start codon (top left); the blue target mRNAs contain longer 5' UTR with complicated secondary structures and the Kozak sequence with much less consensus around the start codon (top right). R (IUPAC nucleotide code) indicates A or G, the -3 position of the Kozak sequence (RccAUGG). Representative red and blue targets with predicted structures are shown (middle and bottom). Predicted RNA structures of Rps26 and eIF3K (as examples of red targets), Mettl5 (as an example of blue targets), and Sox2 (as an example of pluripotency targets) are shown. The structures are predicted by the RNAfold webserver (in each box: the left is the result of minimum free energy predicted secondary structure; the right is the result of centroid predicted secondary structure).

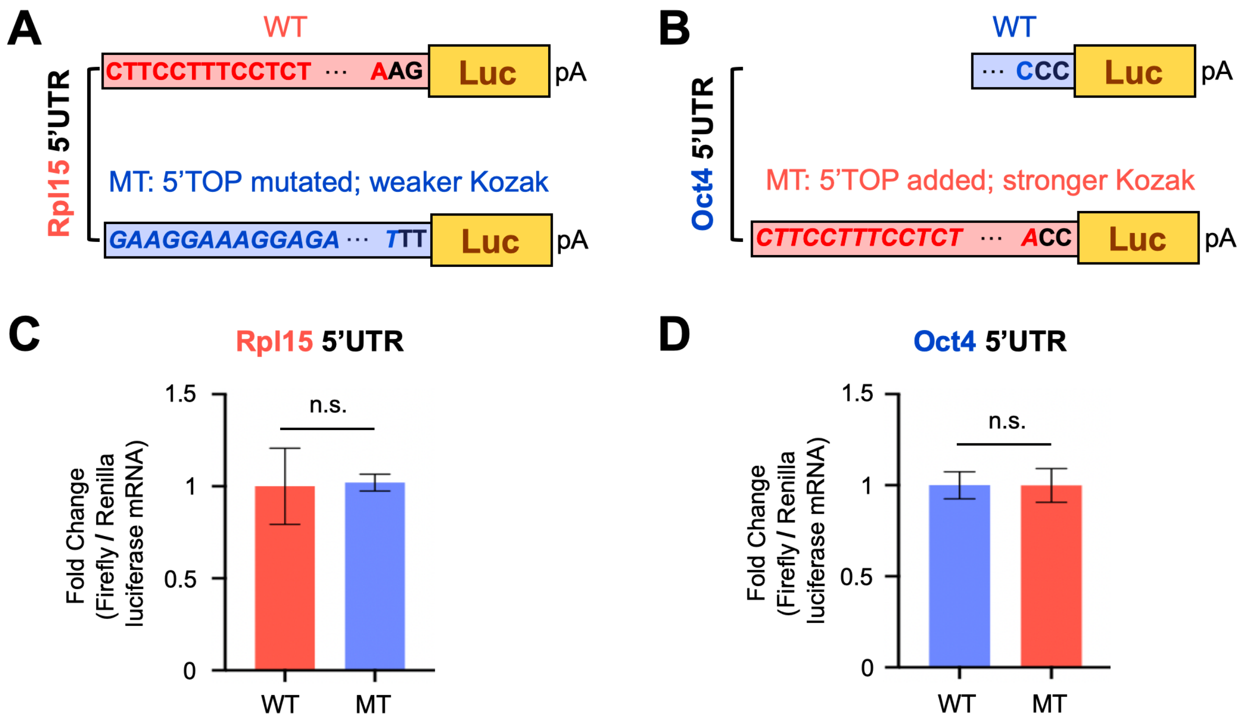
A
WT
Rpl15 5'UTR
CTTCCTTTCCTCT ··· AAG
Luc
pA
MT: 5'TOP mutated; weaker Kozak
GAAGGAAAGGAGA ··· TTT
Luc
pA
B
WT
Oct4 5'UTR
··· CCC
Luc
pA
MT: 5'TOP added; stronger Kozak
CTTCCTTTCCTCT ··· ACC
Luc
pA
C
Rpl15 5'UTR
Fold Change (Firefly / Renilla luciferase mRNA)
1.5
1
0.5
0
n.s.
WT
MT
D
Oct4 5'UTR
Fold Change (Firefly / Renilla luciferase mRNA)
1.5
1
0.5
0
n.s.
WT
MT

**Fig. S4. The 5'TOP motif and the Kozak sequence are important for the Rps26-dependent translational control.**

**A**,**B**, The RNA constructs of the luciferase reporters used in Fig. 4I (A) and 4J (B). **C**,**D**, The qPCR result of the constructs in A (C) and B (D) in the ESCs with shNT and EV plasmid, relative to the expression levels in WT.

**A**

| | GO Term | Genes | Count | *P* value |
|---|---|---|---|---|
| 1 | Single fertilization | H3F3A, H3F3B, CLGN, MFGE8 | 4 | 9.73E-04 |
| 2 | Regulation of centromere complex assembly | H3F3A, H3F3B | 2 | 4.75E-03 |
| 3 | Negative regulation of chromosome condensation | H3F3A, H3F3B | 2 | 4.75E-03 |
| 4 | Telomeric heterochromatin assembly | H3F3A, H3F3B | 2 | 4.75E-03 |
| 5 | Pericentric heterochromatin assembly | H3F3A, H3F3B | 2 | 9.48E-03 |
| 6 | Translation | RPS26, RPL15, EIF3K, MRPS21, SLC25A39 | 5 | 1.49E-02 |
| 7 | Spermatogenesis | H3F3A, H3F3B, CCNB1, CLGN, CADM1 | 5 | 1.56E-02 |
| 8 | Male gonad development | H3F3A, H3F3B, UBB | 3 | 2.91E-02 |
| 9 | Protein folding | TMX1, P3H1, CLGN | 3 | 3.72E-02 |
| 10 | Muscle cell differentiation | H3F3A, H3F3B | 2 | 3.97E-02 |

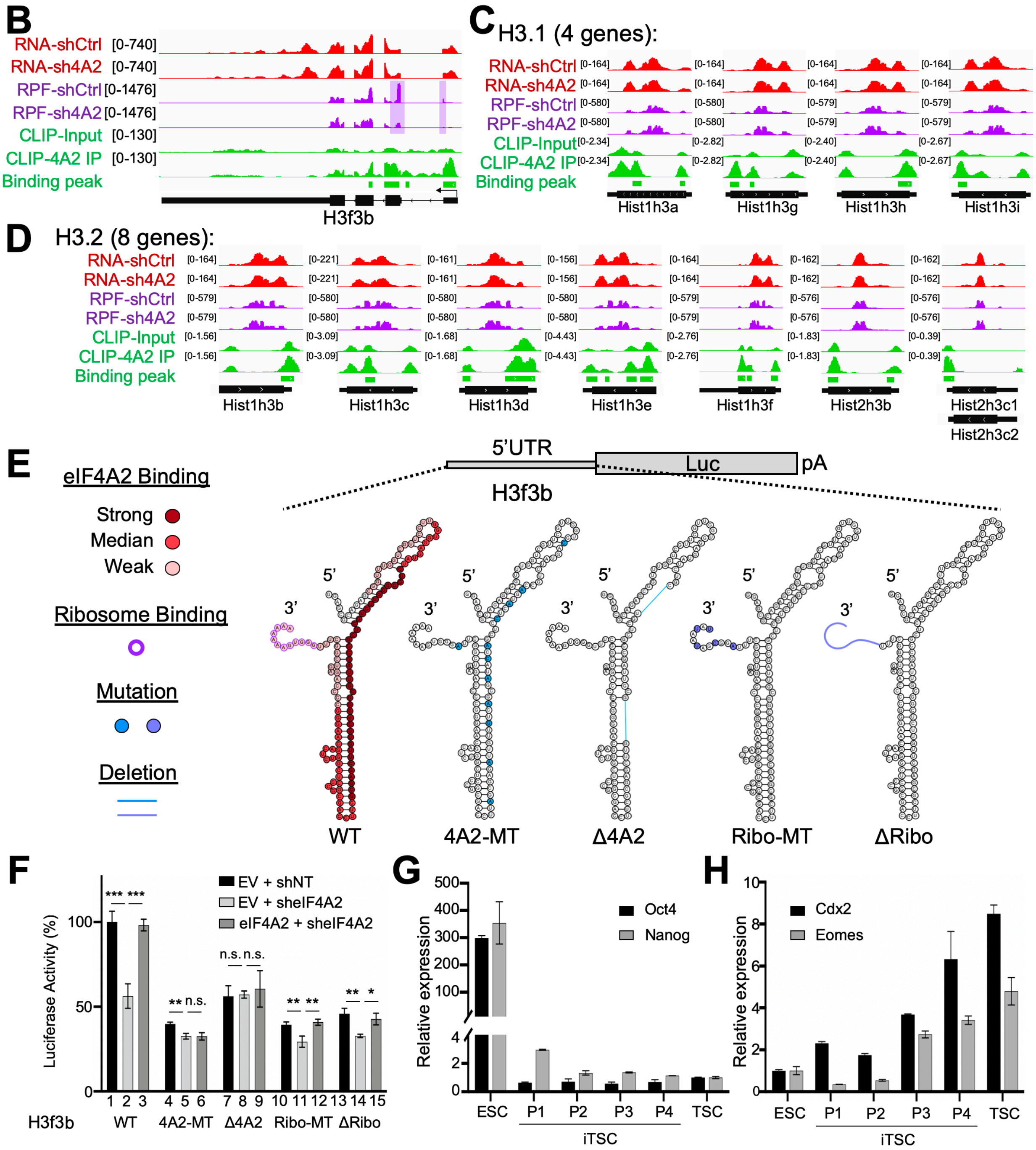

**Fig. S5. eIF4A2 depletion doesn't affect the gene expression of H3.1 or H3.2.**

**A,** The top 10 enriched terms in the GO analysis of the red targets. H3F3A and H3F3B are highlighted in red. **B**,**C**,**D**, IGV snapshots on H3f3b (B), 4 paralogous H3.1 genes (C), and 8 paralogous H3.2 genes (D) with the same datasets as shown in Fig. 2H. **E**, (Top) Luciferase reporter constructs in which the luciferase gene is driven by a H3f3b 5' UTR WT or mutants. (Bottom) The secondary structure of the H3f3b 5' UTR WT or mutants. The indications are the same as the ones in Fig. 5E. **F**, Luciferase activity of mRNAs driven by the H3f3b 5' UTR WT or mutants in ESCs with the indicated shRNA and plasmid. **G**,**H**, The qRT-PCR of (G) pluripotency markers Oct4, Nanog, and (H) TSC markers Cdx2, Eomes, during TSC induction from ZHBTc4 ESCs, relative to the expression levels in ESC, normalized to β-actin expression.

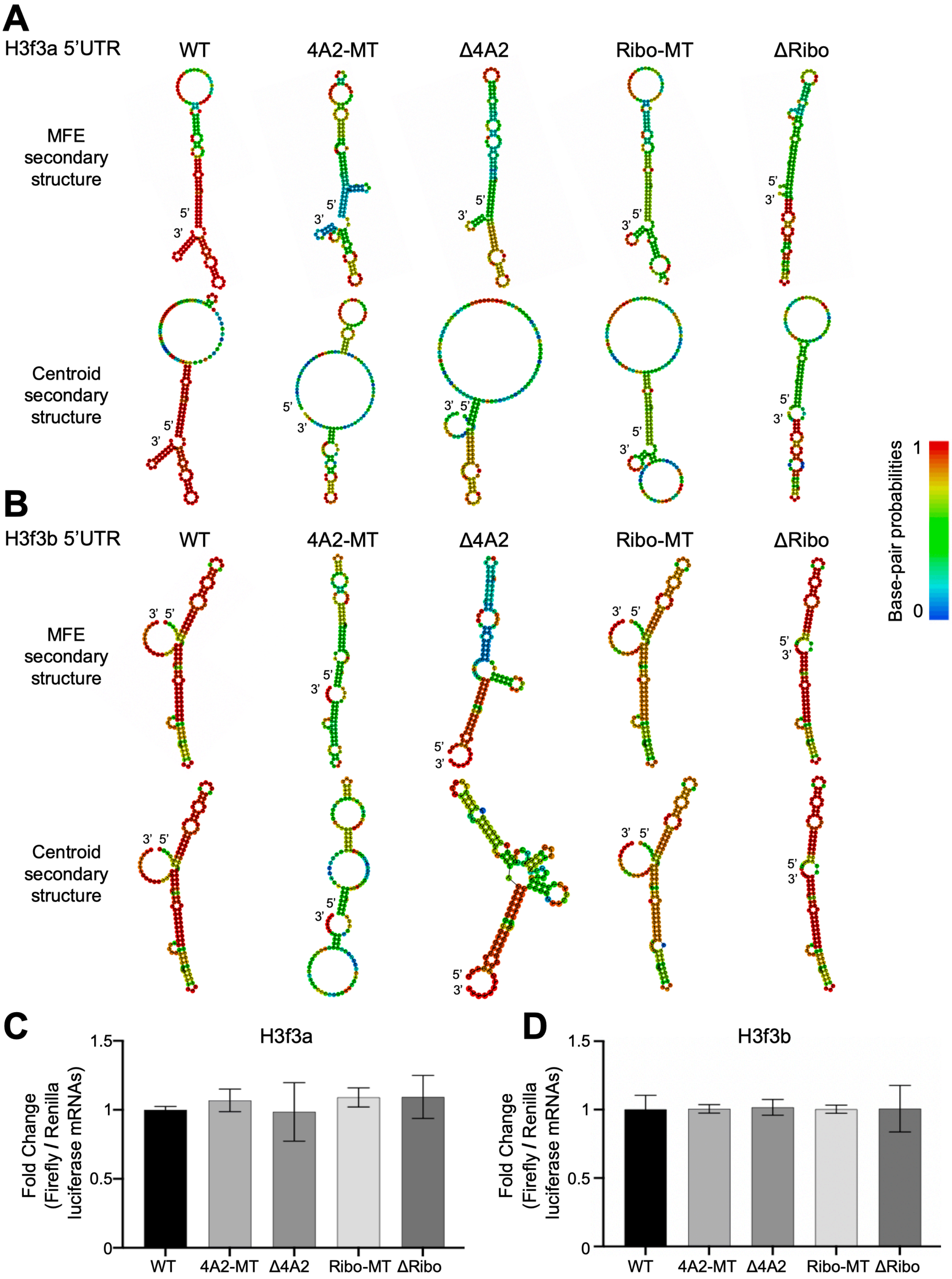

A
H3f3a 5'UTR
WT
4A2-MT
Δ4A2
Ribo-MT
ΔRibo
MFE secondary structure
Centroid secondary structure
B
H3f3b 5'UTR
WT
4A2-MT
Δ4A2
Ribo-MT
ΔRibo
MFE secondary structure
Centroid secondary structure
Base-pair probabilities
1
0
C
H3f3a
Fold Change (Firefly / Renilla luciferase mRNAs)
1.5
1
0.5
0
WT
4A2-MT
Δ4A2
Ribo-MT
ΔRibo
D
H3f3b
Fold Change (Firefly / Renilla luciferase mRNAs)
1.5
1
0.5
0
WT
4A2-MT
Δ4A2
Ribo-MT
ΔRibo

**Fig. S6. The mutations in H3.3 5'UTR reporters don't affect the RNA stability.**

**A,B**, Predicted RNA secondary structures of (A) the H3f3a 5' UTR WT and mutants as well as (B) the H3f3b WT and mutants. The structures are predicted by RNAfold webserver. **C**,**D**, The qPCR result of the constructs in A (C) and B (D) in the ESCs with shNT and EV plasmid, relative to the expression levels in WT.

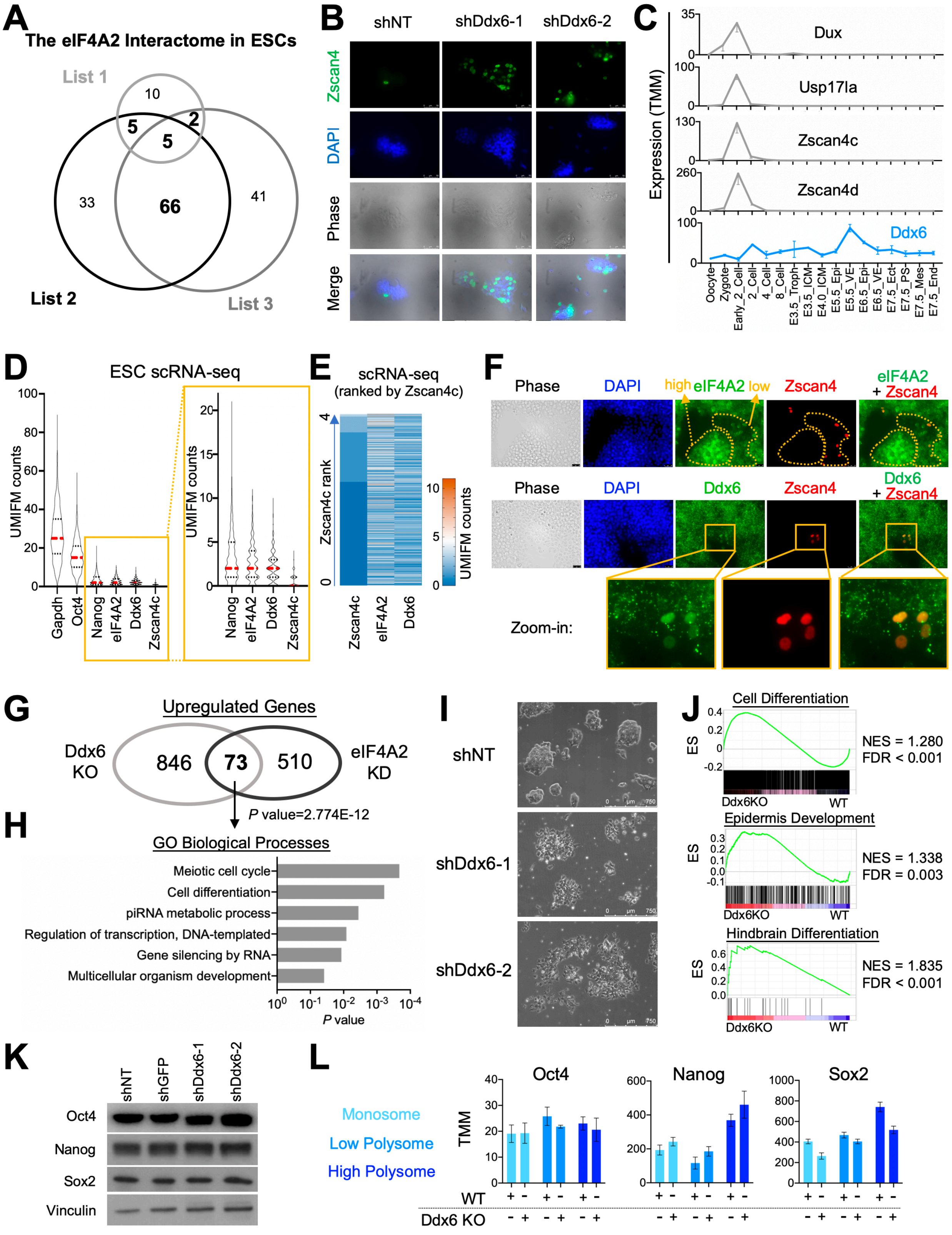

A
The eIF4A2 Interactome in ESCs
List 1
List 2
List 3
10
5
2
5
33
66
41
B
shNT
shDdx6-1
shDdx6-2
Zscan4
DAPI
Phase
Merge
C
Expression (TMM)
Dux
Usp17la
Zscan4c
Zscan4d
Ddx6
Oocyte
Zygote
Early_2_Cell
2_Cell
4_Cell
8_Cell
E3.5_Troph
E3.5_ICM
E4.0_ICM
E5.5_Epi
E5.5_VE
E6.5_Epi
E6.5_VE
E7.5_Ect
E7.5_PS
E7.5_Mes
E7.5_End
D
ESC scRNA-seq
UMIFM counts
Gapdh
Oct4
Nanog
eIF4A2
Ddx6
Zscan4c
E
scRNA-seq
(ranked by Zscan4c)
Zscan4c rank
UMIFM counts
F
Phase
DAPI
high eIF4A2 low
Zscan4
eIF4A2
+ Zscan4
Ddx6
Ddx6
+ Zscan4
Zoom-in:
G
Upregulated Genes
Ddx6
KO
846
73
510
eIF4A2
KD
P value=2.774E-12
H
GO Biological Processes
Meiotic cell cycle
Cell differentiation
piRNA metabolic process
Regulation of transcription, DNA-templated
Gene silencing by RNA
Multicellular organism development
P value
I
shNT
shDdx6-1
shDdx6-2
J
Cell Differentiation
ES
Ddx6KO
WT
NES = 1.280
FDR < 0.001
Epidermis Development
NES = 1.338
FDR = 0.003
Hindbrain Differentiation
NES = 1.835
FDR < 0.001
K
shNT
shGFP
shDdx6-1
shDdx6-2
Oct4
Nanog
Sox2
Vinculin
L
Monosome
Low Polysome
High Polysome
Oct4
Nanog
Sox2
TMM
WT
Ddx6 KO

**Fig. S7. eIF4A2 and Ddx6 inhibit expression of Zscan4 in ESCs.**

**A,** Venn diagrams showing the overlap of the protein interactors identified in three eIF4A2 IP-MS in ESCs. **B**, Immunofluorescence of endogenous Zscan4 in ESCs with control shRNA (shNT) or Ddx6 shRNAs. **C**, The graph showing RNA expression levels of 2C genes and Ddx6 during mouse embryogenesis. Data is from GSE76505 and GSE66390. **D**, The UMIFM counts of the indicated RNA from the ESC scRNA-seq. The zoom-in on the right is to show the UMIFM counts of Nanog, eIF4A2, Ddx6, and Zscan4c. **E**, The heatmap of the UMIFM counts of Zscan4c, eIF4A2, and Ddx6 from the ESC scRNA-seq, ranked by the counts of Zscan4c from 0 (lowest) to 4 (highest). **F**, Immunofluorescence of Zscan4 and eIF4A2 (top), Zscan4 and Ddx6 (middle) in ESCs. The yellow dashed lines (top) highlight the regions with eIF4A2 high (left) or low (right) expression. The zoom-in (bottom) shows the Zscan4+ cell region in the co-staining of Zscan4 and Ddx6 (middle). **G**,**H**, Venn diagrams showing upregulated transcripts upon Ddx6 KO or eIF4A2 KD in ESCs (G) with corresponding GO terms of overlapped genes (H). **I,** Cell morphology of ESCs with either control shRNA (shNT) or Ddx6 shRNAs. **J**, GSEA analyses of the indicated gene sets by comparing Ddx6 KO with WT cells from the RNA-seq data. NES, normalized enrichment score. **K**, Western blots of Oct4, Nanog, and Sox2 in ESCs with control KD and Ddx6 KD. Vinculin serves as the loading control. **L**, Polysome profiling data of Oct4, Nanog, and Sox2 in WT or Ddx6 KO ESCs.
